# Phylogroup-specific variation shapes the clustering of antimicrobial resistance genes and defence systems across regions of genome plasticity

**DOI:** 10.1101/2022.04.24.489302

**Authors:** João Botelho, Leif Tüffers, Janina Fuss, Florian Buchholz, Christian Utpatel, Jens Klockgether, Stefan Niemann, Burkhard Tümmler, Hinrich Schulenburg

**Affiliations:** Antibiotic resistance group, Max-Planck Institute for Evolutionary Biology, Plön, Germany; Evolutionary Ecology and Genetics, University of Kiel, Kiel, Germany; Department of Infectious Diseases and Microbiology, University of Lübeck, Lübeck, Germany; Institute of Clinical Molecular Biology, Christian Albrechts University and University Hospital Schleswig-Holstein, Kiel, Germany; Molecular and Experimental Mycobacteriology, Research Center Borstel, Borstel, Germany; German Center for Infection Research, Partner Site Hamburg-Lübeck-Borstel-Riems, Borstel, Germany; Clinic for Paediatric Pneumology, Allergology, and Neonatology, Hannover Medical School (MHH), Hannover, Germany; Biomedical Research in Endstage and Obstructive Lung Disease Hannover (BREATH), German Center for Lung Research, Hannover Medical School, Hannover, Germany

**Author notes:** To whom correspondence should be addressed. Correspondence may also be addressed to.

## Abstract

**Background:** *Pseudomonas aeruginosa* is an opportunistic pathogen consisting of three phylogroups (hereafter named A, B, and C) of unevenly distributed size. Here, we assessed phylogroup-specific evolutionary dynamics in a collection of *P. aeruginosa* genomes.

**Methods:** In this genomic analysis, using phylogenomic and comparative genomic analyses, we generated 18 hybrid assemblies from a phylogenetically diverse collection of clinical and environmental *P. aeruginosa* isolates, and contextualised this information with 1991 publicly available genomes of the same species. We explored to what extent antimicrobial resistance (AMR) genes, defence systems, and virulence genes vary in their distribution across regions of genome plasticity (RGPs) and “masked” (RGP-free) genomes, and to what extent this variation differs among the phylogroups.

**Findings:** We found that members of phylogroup B possess larger genomes, contribute a comparatively larger number of pangenome families, and show lower abundance of CRISPR-Cas systems. Furthermore, AMR and defence systems are pervasive in RGPs and integrative and conjugative/mobilizable elements (ICEs/IMEs) from phylogroups A and B, and the abundance of these cargo genes is often significantly correlated. Moreover, inter- and intra-phylogroup interactions occur at the accessory genome level, suggesting frequent recombination events. Finally, we provide here a panel of diverse *P. aeruginosa* strains to be used as reference for functional analyses.

**Interpretation:** Altogether, our results highlight distinct pangenome characteristics of the *P. aeruginosa* phylogroups, which are possibly influenced by variation in the abundance of CRISPR-Cas systems and that are shaped by the differential distribution of other defence systems and AMR genes.

**Funding:** German Science Foundation, Max-Planck Society, Leibniz ScienceCampus Evolutionary Medicine of the Lung, BMBF program Medical Infection Genomics, Kiel Life Science Postdoc Award.

**Research in context:** *Evidence before this study:* To date, pangenome studies exploring the epidemiology and evolution dynamics of bacterial pathogens have been limited due to the use of gene frequencies across whole species dataset without accounting for biased sampling or the population structure of the genomes in the dataset. We searched PubMed without language restrictions for articles published before September 1, 2021, that investigated the phylogroup-specific evolutionary dynamics across bacterial species. In this literature search we used the search terms “pangenome” and “phylogroup” or “uneven”, which returned 14 results. Of these, only one study used a population structure-aware approach to explore pangenome dynamics in a bacterial species consisting of multiple phylogroups with unevenly distributed members.

*Added value of this study:* To our knowledge, this study is the first to assess phylogroup-specific evolutionary dynamics in a collection of genomes belonging to the nosocomial pathogen *P. aeruginosa.* Using a refined approach that challenges traditional pangenome analyses, we found specific signatures for each of the three phylogroups, and we demonstrate that members of phylogroup B contribute a comparatively larger number of pangenome families, have larger genomes, and have a lower prevalence of CRISPR-Cas systems. Additionally, we observed that antibiotic resistance and defence systems are pervasive in regions of genome plasticity and integrative and conjugative/mobilizable elements from phylogroups A and B, and that antibiotic resistance and defence systems are often significantly correlated in these mobile genetic elements.

*Implications of all the available evidence:* These results indicate that biases inherent to traditional pangenome approaches can obscure the real distribution of important cargo genes in a bacterial species with a complex population structure. Furthermore, our findings pave the way to new pangenome approaches that are currently under-explored in comparative genomics and, crucially, shed a new light on the role that integrative and conjugative/mobilizable elements may play in protecting the host against foreign DNA.

## Introduction

*Pseudomonas aeruginosa* is a ubiquitous metabolically versatile γ-proteobacterium. This Gram-negative bacterium is also an opportunistic human pathogen commonly linked to life-threatening acute and chronic infections ^1^. It belongs to the ESKAPE pathogens collection ^2^, highlighting its major contribution to nosocomial infections across the globe and its ability to “escape” antimicrobial therapy because of the widespread evolution of antimicrobial resistance (AMR) ^3^. This species is also often found to be multi-as well as extensively drug resistant (MDR and XDR, respectively) ^4^, making it difficult and in some cases even impossible to treat. For this reason, *P. aeruginosa* is placed by the World Health Organization (WHO) in the top priority group of most critical human pathogens, for which new treatment options are urgently required ^5^. These efforts rely on an in-depth understanding of the species biology and its evolutionary potential, which may be improved through a functional analysis of whole genome sequencing data.

The combined pool of genes belonging to the same bacterial species is commonly referred to as the pangenome. Frequently, only a small proportion of these genes is shared by all species members (the core genome). On the contrary, a substantial proportion of the total pool of genes is heterogeneously distributed across the members (the accessory genome). Following Koonin and Wolf ^6^, the pangenome can be divided into 3 categories: i) the persistent or softcore genome, for gene families present in the majority of the genomes; ii) the shell genome, for those present at intermediate frequencies and that are gained and lost rather slowly; iii) the cloud genome, for gene families present at low frequency in all genomes and that are rapidly gained and lost ^7^. Clusters of genes that are part of the accessory genome (i.e, the shell and cloud genome) are often located in so-called regions of genome plasticity (RGPs), genomic loci apparently prone to insertion of foreign DNA. By harbouring divergent accessory DNA in different strains, these loci can represent highly variable genomic regions. The shell and cloud genomes are also characterized by mobile genetic elements (MGEs) that are capable of being laterally transferred between bacterial cells, including plasmids, integrative and conjugative/mobilizable elements (ICEs/IMEs), and prophages ^8,9^. These MGEs can mediate the shuffling of cargo genes that may provide a selective advantage to the recipient cell, such as resistance to antibiotics, increased pathogenicity, and defence systems against foreign DNA ^10–12^.

Most pangenome studies described to date have characterized gene frequencies across the whole species dataset without accounting for biased sampling or the population structure of the genomes in the dataset. This is particularly relevant in species consisting of multiple phylogroups with unevenly distributed members. As recently reported for *Escherichia coli* ^13^, genes classified as part of the accessory genome using traditional pangenome approaches are in fact core to specific phylogroups. Since *P. aeruginosa* is composed of three different-sized phylogroups (hereafter referred to as phylogroups A, B, and C as per the nomenclature proposed by Ozer *et al* ^14^; see also results), characterized by high intraspecies functional variability ^15,16^, it is likely that evolution in these phylogroups is driven by specific sets of genes found in the majority of members within the groups, but not across groups.

The aim of the current study is to enhance our understanding of the pangenome of the human pathogen *P. aeruginosa* by specifically assessing phylogroup-specific characteristics and genome dynamics, including data from more than 2000 genomes. We explore to what extent particular groups of cargo genes, such as those encoding AMR, virulence, and defence systems, vary in their distribution across RGPs and “masked” (RGP-free) genomes, and to what extent this variation differs among the phylogroups. Our data set includes new full genome sequences of a representative set of *P. aeruginosa* strains, the ‘major *P. aeruginosa* clone type’ (mPact) strain panel. This set of strains was previously isolated by the Tümmler lab (Hanover, Germany) from both clinical and environmental samples ^17^. This mPact panel encompasses the most common clone types in the contemporary population ^18–20^ and provides a manageable, focused resource for in-depth functional analyses.

## Methods

### Sequencing and hybrid assembly of the mPact strain panel

Genomic DNA from 18 strains of the mPact panel ^17^ were extracted using the Macherey-Nagel NucleoSpin Tissue kit, according to the standard bacteria support protocol from the manufacturer. We used Nanodrop 1000 for DNA quantification and quality control (260/280 and 260/230 ratios), followed by measurements in Qubit for a more precise quantification. The Agilent TapeStation and the FragmentAnalyzer Genomic DNA 50KB kit served to control fragment size. Sequencing libraries were prepared with the Illumina Nextera DNA flex and Pacific Bioscience (PacBio) SMRTbell express template prep kit 2.0. Libraries were sequenced on the Illumina MiSeq at 2×300bp or the PacBio Sequel II, respectively. Illumina reads were verified for quality using FastQC v0.11.9 ^21^ and trimmed with Trim Galore v0.6.6 ^22^, using the paired-end mode with default parameters and a quality Phred score cutoff of 10. Both datasets were then combined using the Unicycler v0.4.8 assembly pipeline ^23^. We used the default normal mode in Unicycler to build the assembly graphs of most strains, except of the mPact strains H02, H14, H15, H18, and H19, where we used the bold mode. The assemblies were visually inspected using the assembly graph tool Bandage v0.8.1 ^24^.

### Bacterial collection

We downloaded a total of 5468 *P. aeruginosa* genomes from RefSeq’s NCBI database using PanACoTA v1.2.0 ^25^. After quality control to remove low-quality assemblies, 2704 were kept and 2764 genomes with more than 100 contigs were discarded (**Table S1**). Next, 713 genomes were discarded by the distance filtering step, using minimum (1e-4) and maximum (0.05) mash distance cut-offs to remove duplicates and misclassified assemblies at the species level ^26^, respectively. This resulted in 1991 publicly available genomes. The 18 genomes sequenced in this study from mPact panel ^17^ passed both filtering steps, resulting in a pruned collection of 2009 genomes in total. Multi-locus sequence typing (MLST) profiles were determined with mlst v2.19.0 (https://github.com/tseemann/mlst).

### Pangenome and phylogenomics

The average nucleotide identity (ANI) between the 2009 genomes was calculated with fastANI v1.33 ^26^. We used the genome sequences to generate a pangenome with the panrgp subcommand of PPanGGOLiN v1.1.136 ^27,28^. We built a softcore-genome alignment (threshold 95%), followed by inference of a maximum likelihood tree with the General Time Reversible model of nucleotide substitution in IQ-TREE v2.1.2 ^29^. To detect recombination events in our collection and account for them in phylogenetic reconstruction, we used ClonalFrameML v1.12 ^30^. Phylogenetic trees were plotted in iTOL v6 (https://itol.embl.de/)^31^ and explored to cluster genomes according to the phylogroup. Due to the sample size difference, we subsequently focused the analysis on each phylogroup separately. Pangenome analysis was performed for each phylogroup, using the panrgp subcommand from PPanGGOLiN. Core and accessory genes were classified across genomes from different phylogroups with a publicly available R script (https://github.com/ghoresh11/twilight)^13^. We used the gene presence/absence output from the whole collection’s pangenome and the grouping of our genomes according to the phylogroup.

### Identification of RGPs and ICEs/IMEs

To mask all the genomes, we used the RGPs coordinates determined by panrgp for each individual genome as input in bedtools maskfasta v2.30.0 ^32^. We extracted the RGP nucleotide sequences with the help of bedtools getfasta. All genomes were annotated with prokka v1.4.6 ^33^. To look for ICEs/IMEs on complete genomes, we used the genbank files created by prokka as input in the standalone-version of ICEfinder ^34^.

### Identification of ICEs and functional categories

We retrieved the annotated proteins for the RGPs and masked genomes across the three phylogroups. We clustered each of the six groups of proteins with MMseqs2 v13.45111 ^35^ and an identity cut-off of 80%. These clustered proteins were scanned for functional categories in eggNOG-mapper v2 ^36^, using the built-in database for clusters of orthologous groups ^37^. We calculated the relative frequency of these categories by dividing the absolute counts for each category by the total number of clustered proteins found in each of the six groups. CRISPR-Cas systems were identified with the help of CRISPRCasTyper v1.2.3 ^38^. AMRFinder v3.10.18 ^39^ served to locate AMR genes and resistance-associated point mutations. Virulence genes were characterized with the pre-downloaded database from VFDB ^40^ (updated on the 12-05-2021 and including 3867 virulence factors) in abricate v1.0.1 (https://github.com/tseemann/abricate). Finally, we searched for defence systems using the protein sequences generated by prokka as input in defense-finder v0.0.11 ^41^, a tool developed to identify known defence systems in prokaryotic genomes, for which at least one experimental evidence of the defence function is available.

### Network-based analysis of RGPs and ICEs/IMEs

As a first step, we calculated the Jaccard Index between the RGPs with the help of BinDash v0.2.1 ^42^ with *k*-mer size equal to 21 bp. In detail, we used the sketch subcommand to reduce multiple sequences into one sketch, followed by the dist subcommand, to estimate distance (and relevant statistics) between RGPs in query sketch and RGPs in target-sketch. The Jaccard Index between ICEs/IMEs was similarly obtained with BinDash. We used the mean() function in R to calculate the arithmetic mean of the Jaccard Index. Only Jaccard Index values equal to or above the mean were considered, and the mutation distances served as edge attributes to plot the networks with Cytoscape v3.9.1 under the prefuse force directed layout (https://cytoscape.org/). Based on the Analyzer function in Cytoscape, we computed a comprehensive set of topological parameters, such as the clustering coefficient, the network density, the centralization, and the heterogeneity. Clusters in our networks were identified with the AutAnnotate and clusterMaker apps available in Cytoscape, using the connected components as the clustering algorithm.

### Statistical analysis

The correlation matrix was ordered using the hclust function in R. Statistical comparison of the variation between groups was always based on non-parametric tests, thereby taking into account that the compared groups varied in data distributions (e.g., at least one group with a skewed distribution) and/or showed unequal variances. Moreover, as non-parametric tests are usually considered to be conservative, the thus identified significant test results should indicate trustworthy differences between groups. In particular, the three phylogroups (e.g., genome size, GC content) were generally compared using the Kruskal-Wallis test. The unpaired two-sample Wilcoxon test was used in multiple comparisons between two independent groups of samples (RGPs vs. masked genomes, CRISPR-Cas positive vs negative genomes). In both tests, p-values were adjusted using the Holm–Bonferroni method. Values above 0.05 were considered as nonsignificant (ns). We used the following convention for symbols indicating statistical significance: * for p <= 0.05, ** for p <= 0.01, *** for p <= 0.001, and **** for p <= 0.0001.

### Role of funding source

The funders (Max-Planck Society, German Science Foundation, German ministry for education and research, Kiel university) had no role in the study design, data collection, data analysis, data interpretation, or writing of the report. The corresponding authors had full access to all the data and final responsibility for the decision to submit for publication. None of the contributing authors were precluded from accessing data of this study.

## Results

### The *P. aeruginosa* phylogeny is composed of three phylogroups

Our phylogenomic characterization was based on 2009 assembled *P. aeruginosa* genomes belonging to 519 MLST profiles and including 1991 publicly available genomes (following quality control and distance filtering, **Table S2**) and additionally 18 genomes for the mPact strain panel (**Table S3**) ^17^. Analysis of the ANI values (**Figure S1**) and the softcore-genome alignment of these genomes identified three phylogroups, as previously reported ^14,43^ (**Figure 1**). The two major reference isolates are part of the larger phylogroups: PAO1 ^44^ is part of phylogroup A (n=1531), while the PA14 strain falls into phylogroup B (n=435). Phylogroup C includes a substantially smaller number of members (n=43) (**Table S2**). Members of the phylogroup C were recently subdivided into either 2 ^14^ or 3 clusters, including the distantly related PA7 cluster ^43^. In this work, however, the PA7 cluster was excluded, and we focused our analysis on only the remainder of phylogroup C, since genomes from the PA7 cluster were too distantly related to the other genomes. In fact, the PA7 strain was first described as a taxonomic outlier of this species ^45^, and genomes belonging to this cluster were recently proposed to belong to a new *Pseudomonas* species ^46^. To test the impact of recombination on the softcore-genome alignment, we used ClonalFrameML to reconstruct the phylogenomic tree with corrected branch lengths. The segregation of *P. aeruginosa* into three phylogroups was maintained, resulting in a tree with decreased branch lengths and with identical number of members assigned to each phylogroup (**Figure S2**). Genomes from the mPact panel sequenced in this study were widely distributed across the *P. aeruginosa* phylogeny, with 12 strains in phylogroup A, 5 in phylogroup B, and 1 in phylogroup C (**Figure 1** and **Table S3**). Our results show that *P. aeruginosa* consists of three asymmetrical phylogroups and that the segregation of the 2009 genomes into phylogenetically distinct groups is not an artefact of recombination.

**Figure 1.**
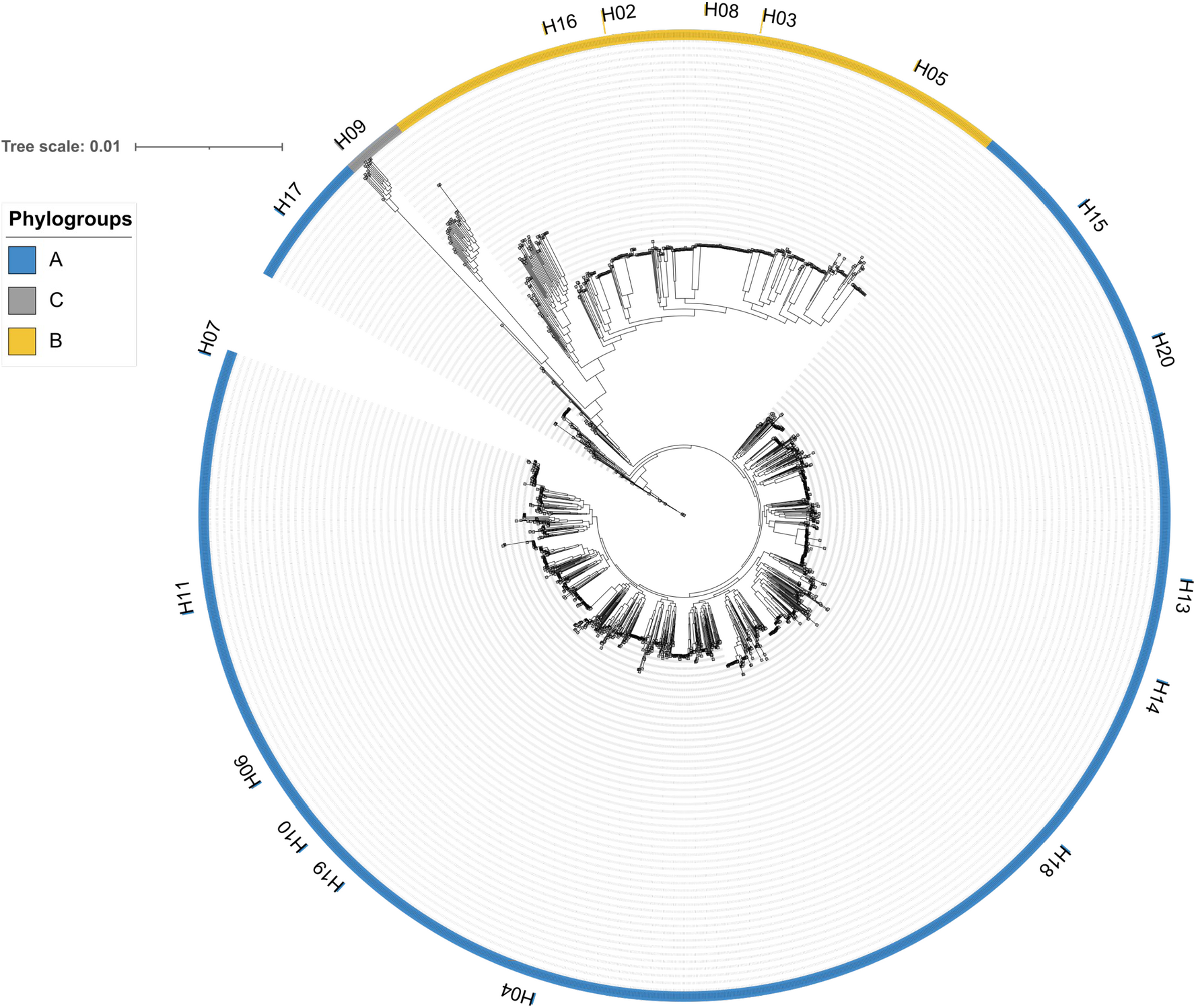
Maximum-likelihood tree of the softcore-genome alignment of all *P. aeruginosa* isolates used in this study (n=2009). The scale bar represents the genetic distance. Arcs in blue represent phylogroup A, yellow B, and grey C. The phylogenetic placement of the major *P. aeruginosa* clone type (mPact) strain panel, sequenced in this study, are highlighted in the tree, with the strain name (the “H” before each number stands for Hanover, referring to the location of the Tuemmler lab and the study that first described this collection ^17^) next to strips coloured according to the phylogroup.

### Phylogroup B contributes comparatively more gene families to the pangenome than the other two phylogroups

We next built a pangenome for the whole species, and separate pangenomes for each of the three phylogroups. This latter approach is important to take phylogenetic subdivisions of the species into account, which is additionally critical because the three phylogroups in our collection have substantially different sample sizes. We observed that the number of persistent gene families in the larger phylogroups A and B were similar to those found in the whole species, while the phylogroup C contained a substantially smaller number of persistent gene families (**Table S4**).

The pangenome of bacterial species is usually classified in two types: open pangenomes and closed ones ^47^. Since *P. aeruginosa* is an example of a bacterial species with open pangenome ^14^, i.e., the sequencing of new genomes will increase pangenome size, we explored the contribution of each phylogroup to the pangenome. To ensure comparability among the three phylogroups in our first analysis, we randomly drew 43 genomes from each phylogroup (thus, including the total sample size of the smallest phylogroup C), and observed that there is more diversity in the accessory genes of phylogroup B as to the functions contributed by the acquired genes (**Figure 2A** and **Table S5**). In our second analysis, we focused only on the two larger phylogroups A and B, for which we randomly drew in each case 100 genomes and found the trend unchanged (**Figure 2B** and **Table S6**).

**Figure 2.**
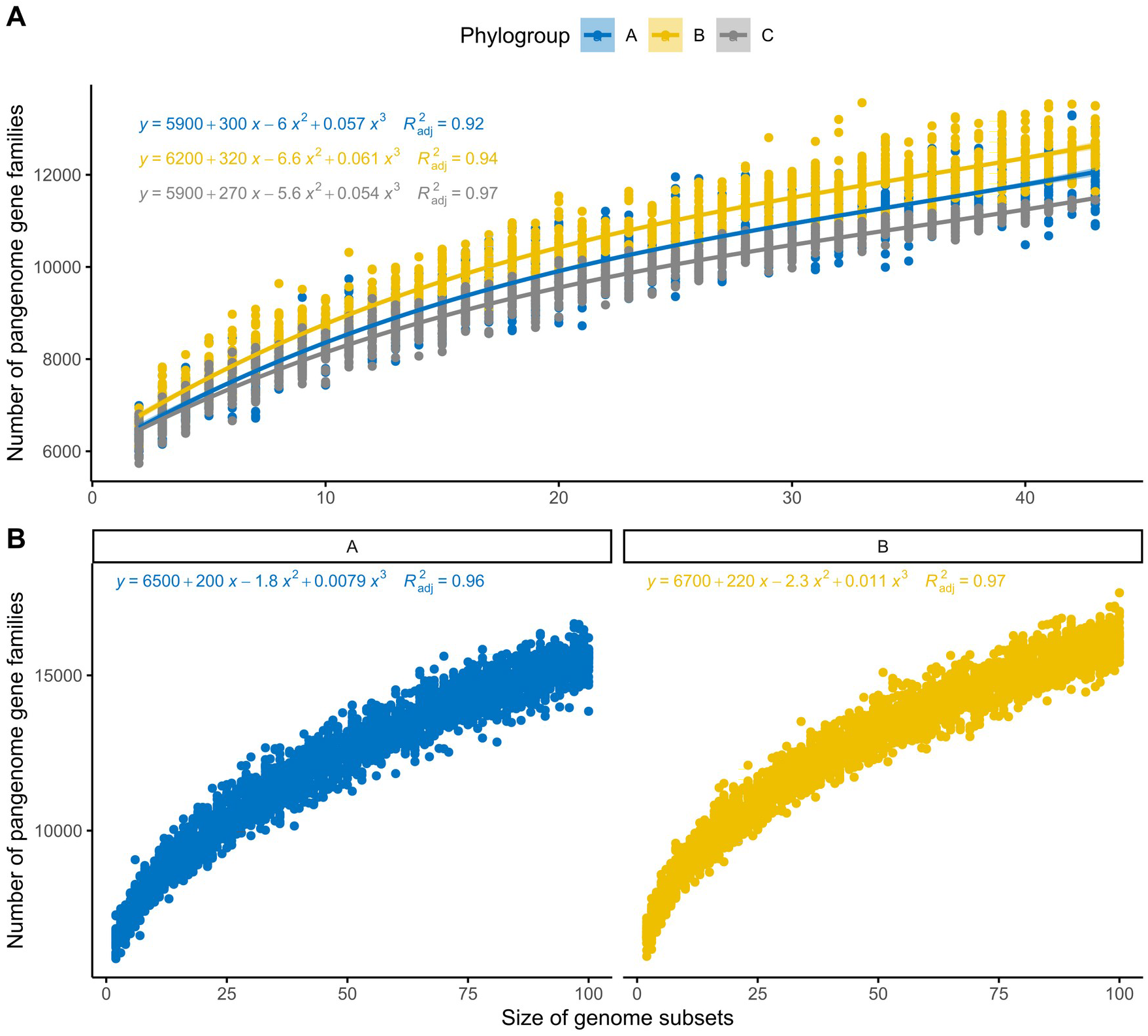
Rarefaction curves of the pangenome gene families for each phylogroup. All curves were inferred using polynomial regression lines. Curves in blue represent phylogroup A, yellow B, and grey C. **A)** The curves were generated by randomly re-sampling 43 genomes from each phylogroup several times and then plotting the average number of pangenome families found on each genome. **B)** Rarefaction curves were plotted with 100 random genomes from phylogroups A and B.

We then explored if specific gene families were pervasive across single or multiple phylogroups. We found 14 phylogroup-specific softcore gene families in phylogroup C, and one gene family each was exclusively found in the softcore genomes of phylogroups A and B, respectively (**Figure S3** and **Table S7**). Most gene families uniquely found on the softcore genome of phylogroup C were part of the Xcp type-II secretion system (T2SS), which is one of two complete and functionally distinct T2SS present in this species (**Table S8**). The Xcp system is encoded in a cluster containing 11 genes (*xcpP–Z),* as well as an additional *xcpA/pilD* gene found elsewhere in the genome ^48^. These genes were also found in the majority of the genomes from phylogroups A and B (**Table S9**), but the encoded proteins were too distantly related to those from phylogroup C. A similar pattern was observed for the two gene families indicated exclusively for either phylogroup A or B, for which we also found distantly related orthologues in phylogroup C. Altogether, these results highlight that phylogroup B differs from the other two by contributing a comparatively larger number of gene families to the pangenome, possibly suggesting that phylogroup B members have larger genomes.

### Phylogroup B genomes are significantly larger and most carry no CRISPR-Cas systems

A comparison of genome lengths revealed significantly larger genome sizes for phylogroup B than the other two phylogroups (**Figure 3A**, p-value < 2.2e-16). We then extracted the RGPs from each phylogroup, and found a total of 57901 RGPs across the three phylogroups. The RGPs from phylogroup B were significantly larger than that of phylogroup A (**Figure S4**), thus at least contributing to the overall size difference. Nevertheless, after removing the RGPs, the resulting “masked” genomes from phylogroup B were still significantly larger than those from the other two phylogroups (**Figure 3B**, p-value < 2.2e-16). Additionally, we found that genomes from phylogroup B are still significantly larger than those from the other two phylogroups, even if phylogroup sample sizes were adjusted to sample size of the smallest group, phylogroup C (with 43 genomes; **Figure S5**, p-value 3.2e-07). These results point to a potentially higher number of genes conserved across genomes from phylogroup B. Still, the difference in genome size between phylogroups A and B is mainly explained by differences in accessory genome size (**Figure S4**). Masked genomes from phylogroup C are significantly smaller than genomes from the other two phylogroups, which is consistent with the smaller number of persistent gene families identified in this phylogroup (**Table S4**). We further explored variation in GC content and observed that the GC content from phylogroup B genomes is significantly lower than those from other phylogroups (**Figure S6**, p-value < 2.2e-16).

**Figure 3.**
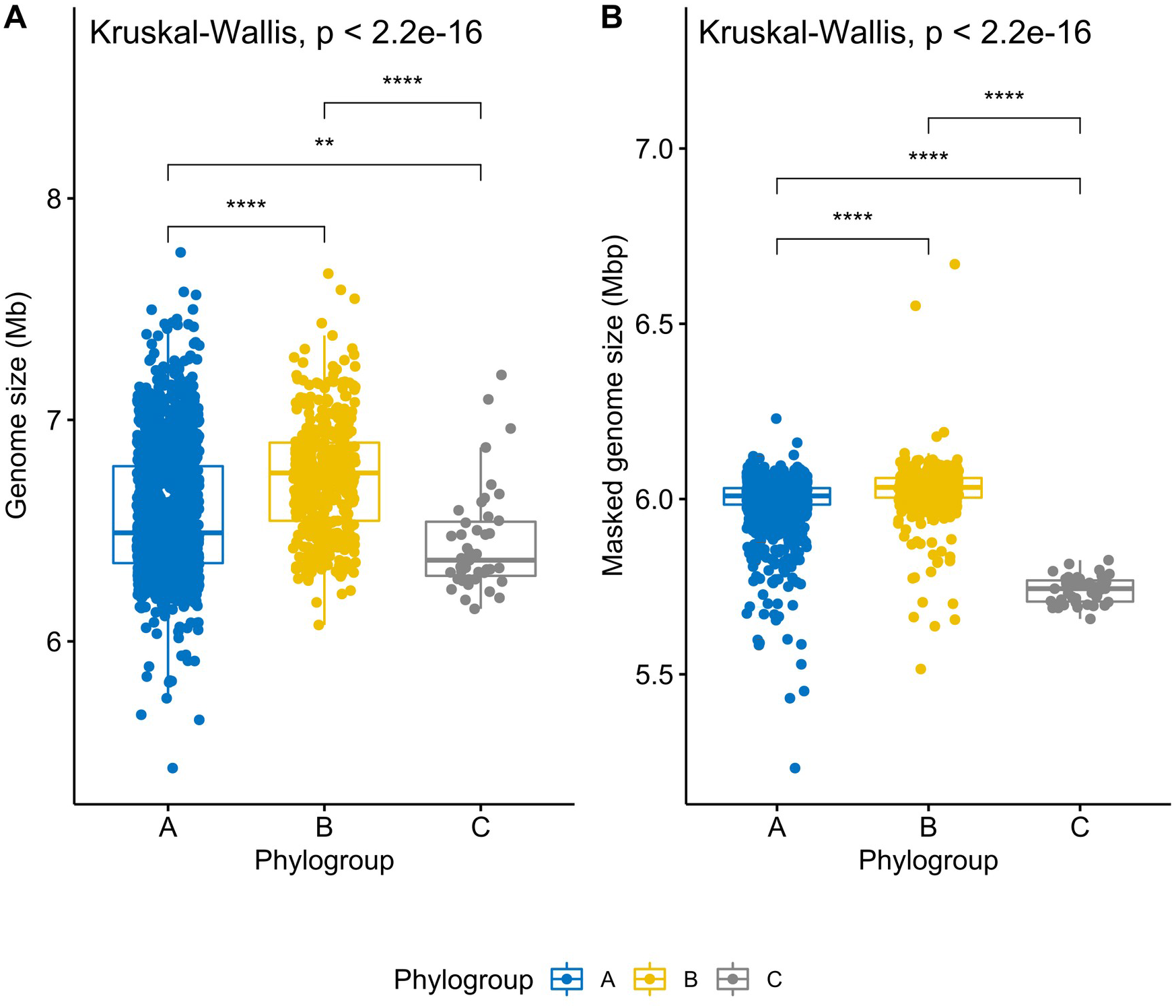
Boxplots representing the variation in genome size **(A)** and masked genome size **(B)** across the three phylogroups. Values above 0.05 were considered as non-significant (ns). Stars indicate significance level: * p <= 0.05, ** p <= 0.01, *** p <= 0.001, and **** p <= 0.0001. Boxplots in blue represent phylogroup A, yellow B, and grey C.

We next assessed whether presence of the defence CRISPR-Cas system is associated with genome size variation. Since CRISPR-Cas systems are important to defend bacteria against foreign DNA ^12,49^, we expected that genomes carrying these systems would be smaller, while those devoid of these systems would accumulate mobile elements and hence be larger. We subdivided genomes from each of the three phylogroups into two groups depending on whether they contain or lack CRISPR-Cas systems, respectively (CRISPR-Cas^pos^, CRISPR-Cas^neg^). We indeed found that genomes with CRISPR-Cas systems are significantly smaller than those without (**Figure 4A**, p-values 8.3e-05 and 0.00025 for the phylogroup A and B comparisons, respectively), supporting the hypothesis that CRISPR-Cas systems can constrain horizontal gene transfer in *P. aeruginosa* ^50–52^. While the number of CRISPR-Cas^pos^ and CRISPR-Cas^neg^ genomes in phylogroups A and C is evenly distributed, phylogroup B genomes without CRISPR-Cas (n=279) were nearly two times more prevalent than those that carried these systems (n=156, **Table S2**). Interestingly, masked genome size of CRISPR-Cas^pos^ and CRISPR-Cas^neg^ phylogroup B isolates was no longer significantly different from one another (**Figure 4B**). In line with this finding, we observed that the cumulative size of all RGPs was higher in genomes without CRISPR-Cas systems across phylogroups A and B (**Figure S7**). The absence of these defence systems in most genomes from phylogroup B may help to explain the observed larger size.

**Figure 4.**
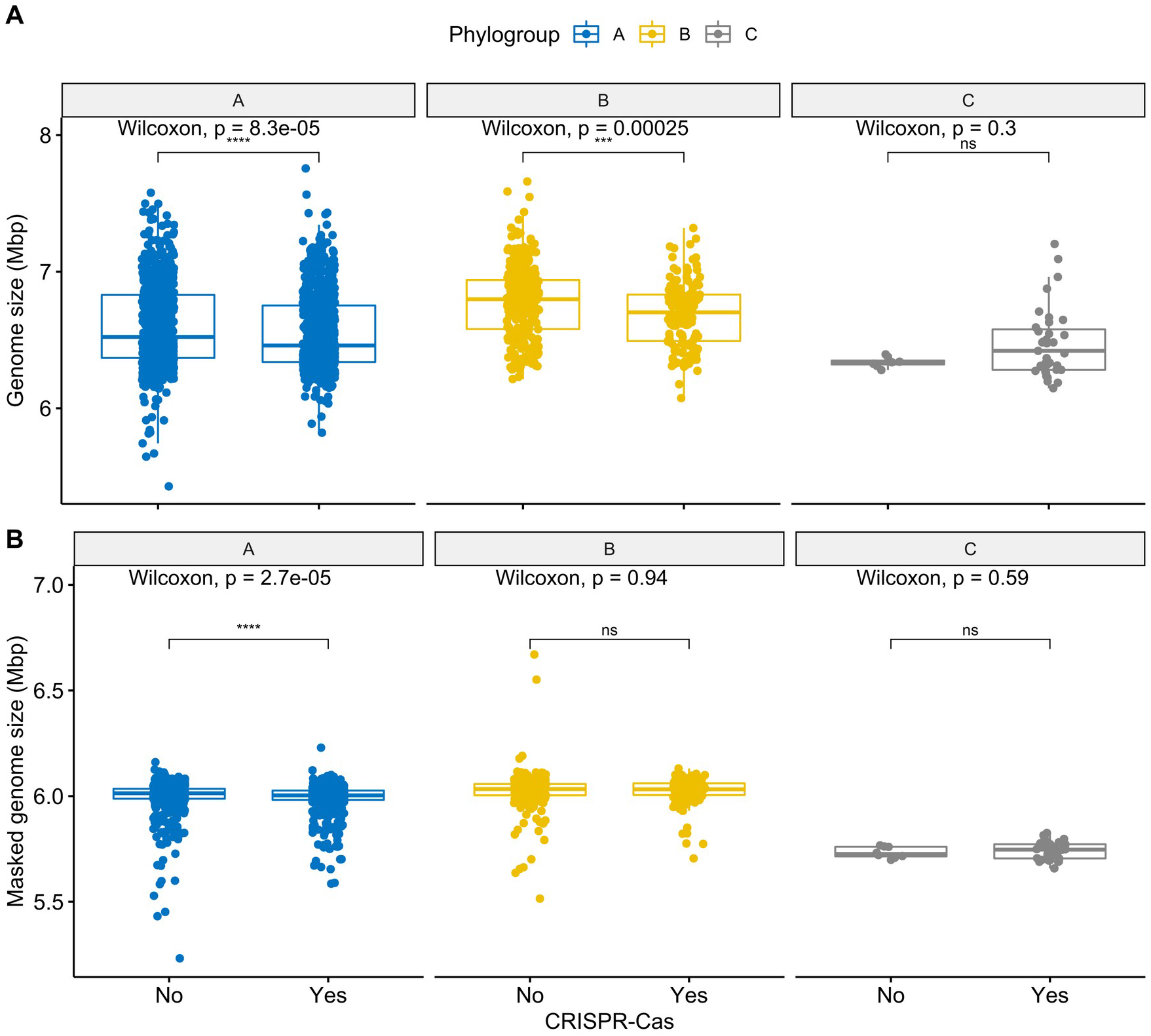
Boxplots representing the variation in genome size **(A)** and masked genome size **(B)** across pairs of conspecific genomes from the same phylogroup with and without CRISPR-Cas systems. Values above 0.05 were considered as non-significant (ns). Stars indicate significance level: * p <= 0.05, ** p <= 0.01, *** p <= 0.001, and **** p <= 0.0001. Boxplots in blue represent phylogroup A, yellow B, and grey C.

We observed a wider diversity of CRISPR-Cas systems in genomes from phylogroup A, including I-C, IE, I-F, IV-A1, and IV-A2 (**Figure S8** and **Table S10**). These CRISPR-Cas subtypes were all found in genomes from phylogroup B, with the exception of the IV-A2. Curiously, only subtypes I-E and I-F were present in phylogroup C. Type IV CRISPR-Cas systems were found almost exclusively on plasmids, and recent work revealed that they participate in plasmid–plasmid warfare ^12,53^. The type I-C CRISPR–Cas subtype is typically encoded on ICEs and is also involved in competition dynamics between mobile elements ^51,54^. Overall, our findings show that phylogroup B genomes are significantly larger and have a wider pool of accessory genes than those from the other two phylogroups, possibly driven by the lower prevalence of CRISPR-Cas systems in phylogroup B.

### AMR and defence systems are overrepresented in RGPs from phylogroups A and B

We next assessed variation in the relative frequency of proteins encoded in RGPs from different phylogroups. We observed that most functional categories are conserved across phylogroups. However, proteins coding for replication, recombination and repair functions are more prevalent in phylogroups A and B RGPs than those from phylogroup C (**Figure 5A**). Since these proteins are frequently involved in mobilization, this finding may suggest that genomes in these phylogroups have more functional mobile elements, with the ability to be horizontally transferred, while the RGPs in phylogroup C may be derived from remnants of mobile elements that can no longer be mobilized.

**Figure 5.**
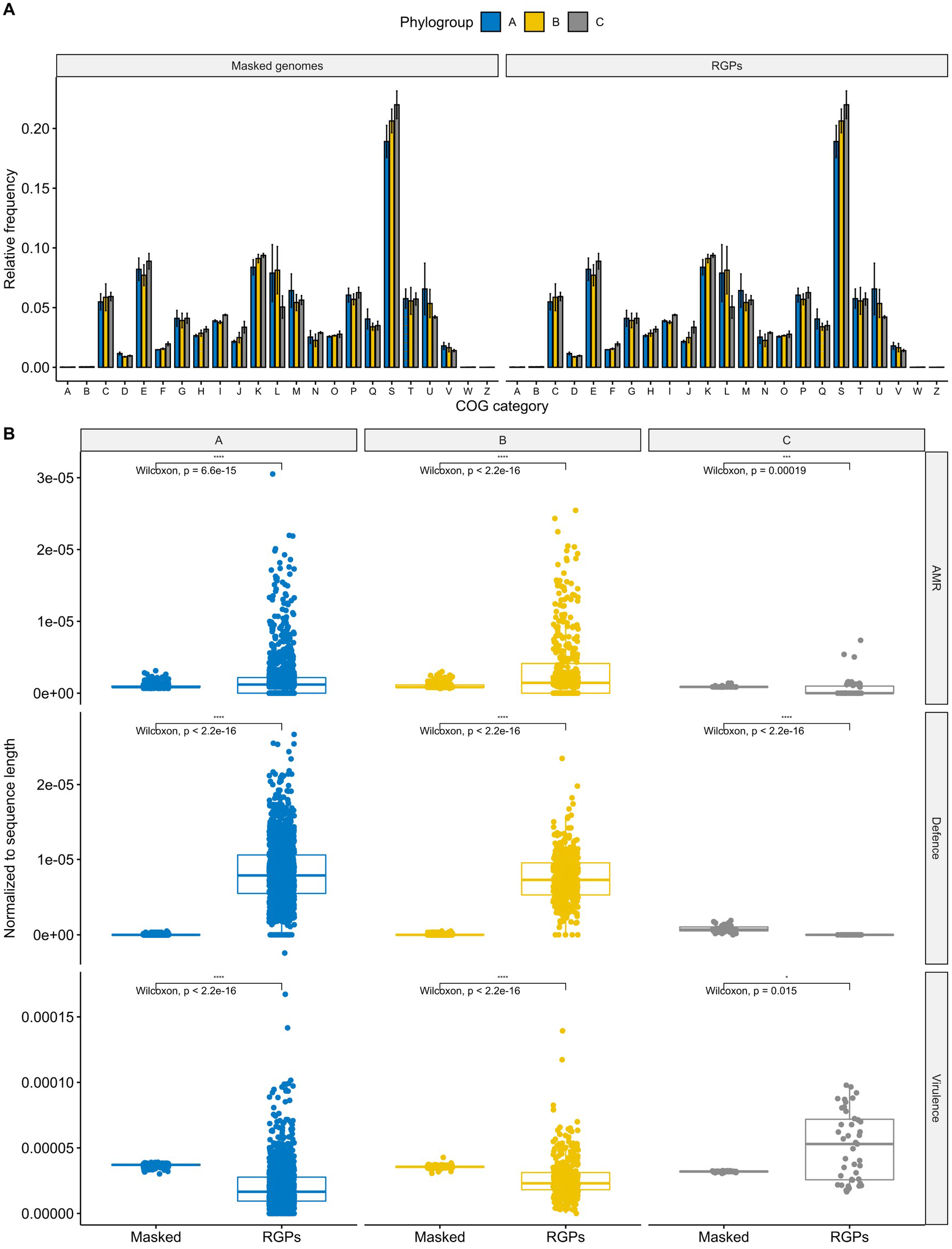
Distribution of functional categories across RGPs and masked genomes from the different phylogroups. Bar and boxplots in blue represent phylogroup A, yellow B, and grey C. **A.** Relative frequencies of cluster of orthologous groups categories. The relative frequencies were calculated by dividing the absolute counts for each category by the total number of clustered proteins found in each of the six groups. Error bars indicate the degree of variation across each COG category from each phylogroup across RGPs and masked genomes. The functional categories are indicated by capital letters, including: A, RNA processing and modification; B, chromatin structure and dynamics; C, energy production and conversion; D, cell cycle control and mitosis; E, amino acid metabolism and transport; F, nucleotide metabolism and transport; G, carbohydrate metabolism and transport; H, coenzyme metabolism; I, lipid metabolism; J, translation; K, transcription; L, replication, recombination and repair; M, cell wall/membrane/envelop biogenesis; N, cell motility; O, post-translational modification, protein turnover, chaperone functions; P, inorganic ion transport and metabolism; Q, secondary structure; R, general functional prediction only; S, function unknown; T, signal transduction; U, intracellular trafficking and secretion; V, defence mechanisms; W, extracellular structures; Z, cytoskeleton. **B** Boxplots of the variation in the number of AMR genes, defence systems, and virulence genes found in RGPs and masked genomes across the three phylogroups. Absolute counts of genes and systems were normalized to RGP and masked genome sequence lengths in each strain. Values above 0.05 were considered as non-significant (ns). Stars indicate significance level: * p <= 0.05, ** p <= 0.01, *** p <= 0.001, and **** p <= 0.0001.

We next assessed to what extent RGPs and masked genomes vary in prevalence of genes for three types of functions, which are often encoded on MGEs, including virulence, defence systems, and AMR. Since the cumulative size of all RGPs is substantially smaller than that of masked genomes (**Table S2**), the number of virulence genes, defence systems, and AMR genes were normalized to the sequence length of the RGPs and masked genomes for each strain. We observed that the gene prevalence for these functions is conserved across masked genomes from different phylogroups, while they are unevenly distributed in RGPs. (**Figure 5B**).

Two important virulence factors were only present in some genomes from phylogroup C, and absent from the other two phylogroups (**Table S9**). These genes *(exlA* and *exlB)* encode hemolysins, and when genomes from phylogroup C carry these genes, the typical type-III secretion system (T3SS) machinery found in most bacteria (encoding the toxins ExoS, ExoY, ExoT, and ExoU) is absent from these genomes, supporting previous reports that these are mutually exclusive ^55^. In agreement with previous findings ^14^, we further found that two important genes encoding T3SS effector proteins *(exoS* and *exoU)* were unevenly distributed across the phylogroups: the *exoS* gene was pervasive among genomes from phylogroup A (99.5%, 1524/1531) and the majority of phylogroup C strains (28/43), while the *exoU* gene was overrepresented in genomes from phylogroup B (408/435) and nearly absent in genomes from the other two phylogroups (**Table S9**). Surprisingly, we also found 23 genomes with the atypical exoS^+^/exoU^+^ genotype, all belonging to phylogroup A (**Table S9**). A high frequency of this genotype has recently been reported in patients from the Brazilian Amazon and Peruvian hospitals ^56,57^. As expected ^58^, some virulence genes were exclusively found on RGPs (i.e., absent from masked genomes): flagellar-associated proteins *fleI/flag, flgL, fliC* and *fliD,* as well as *wzy,* which codes for an O-antigen chain length regulator. All these virulence genes were found in RGPs from both phylogroups A and B.

In agreement with the important role of MGEs as vectors for AMR genes in *P. aeruginosa* ^9,59^, we found that AMR genes were overrepresented in RGPs from phylogroups A and B (**Figures 5B and S9**). We then calculated the relative proportion of different AMR classes across RGPs from the three phylogroups, revealing that most AMR classes were overrepresented across RGPs from phylogroup B (**Figure S10**). This result is consistent with our finding that RGPs play a significant role in the larger genome sizes from this phylogroup (**Figure S4**). Point mutations linked to resistance to beta-lactams and quinolones were observed for all phylogroups (**Table S11**).

A wide array of defence systems with a patchy distribution in closely related and distantly related strains was recently characterized in *P. aeruginosa*, suggesting high rates of horizontal gene transfer ^41^. According to this hypothesis, we would expect to observe an abundance of defence systems in RGPs, when compared with masked genomes. Similar to our results for AMR genes, we found that defence systems are indeed overrepresented in RGPs from phylogroups A and B (**Figure 5B**). Defence systems such as the globally distributed restriction-modification and CRISPR-Cas systems were common in RGPs from both phylogroups. Some rarer systems such as cyclic-oligonucleotide-based anti-phage signalling systems (CBASS) ^60^, Zorya, Gabija, Druantia ^61^, abortive infection ^62^, and bacteriophage exclusion (BREX) ^63^ were also observed in RGPs from phylogroups A and B (**Figure S11** and **Table S12**). In contrast, dGTPases were absent from both phylogroups. Finally, we also observed that AMR and defence systems are overrepresented in specific MLST profiles, including the high-risk clones ST111 and ST233 (**Figure S12**) ^1^. Our results revealed that AMR and defence systems are pervasive in RGPs from phylogroups A and B, and the majority of AMR classes are overrepresented in RGPs from phylogroup B.

### AMR and defence systems are prevalent in ICEs/IMEs from phylogroups A and B

Given that the distribution and clustering of defence systems in *P. aeruginosa* is not dependent on the phylogenetic distance between all strains ^41^, and considering the high prevalence of ICEs/IMEs in this species ^64^, we explored the potential role of these elements as defence islands. To accurately detect these MGEs, we focused our analysis on complete genomes. We noted that 12.6% of our collection consisted of complete genomes (254/2009), including 172 genomes from phylogroup A, 78 from phylogroup B, and 4 genomes from phylogroup C (**Table S2**). 215 out of the 254 complete genomes harboured a total of 477 ICEs and 76 IMEs (**Table S13**). These ICEs/IMEs were present in 136 genomes from phylogroup A, 77 from phylogroup B, and 2 from phylogroup C. Thus, ICEs/IMEs were pervasive in strains from phylogroup B (77/78) and in the majority of strains from phylogroup A (136/172).

Nearly half of the ICEs/IMEs carried at least one AMR gene (228/553), with the ciprofloxacin-modifying *crpP* gene and the sulphonamide-resistance *sul1* gene being most frequent (**Table S14**). Indeed, the *crpP* gene was recently shown to be widely dispersed across ICEs from *P. aeruginosa* ^65^. Around one third of the ICEs/IMEs (193/553) carried at least one defence system, resulting in a total of 250 defence systems across the ICEs/IMEs and including 27 different types (**Figure S13** and **Table S14**). The most frequent defence subtypes were CBASS-III and restriction-modification type-II ^60,62^. Virulence genes were present in a smaller proportion of the ICEs/IMEs (99/553) and showed higher variation in abundance across ICEs/IMEs than AMR genes and defence systems do (**Figure S14**). The *exoU* gene encoding for the effector protein and the *spcU* gene encoding for its chaperone were the most frequent virulence genes, all in ICEs/IMEs from phylogroup B (**Table S14**).

We next explored to what extent the prevalence of these three functional groups is correlated across ICEs/IMEs from the two larger phylogroups A and B. We observed that genes encoding resistance to fluoroquinolones were negatively correlated with genes involved in resistance to other antibiotic classes, and also with specific defence systems as restriction-modification and CBASS (**Figure 6A**). ICEs/IMEs from phylogroup B carrying fluoroquinolone-encoding resistance genes were also negatively associated with genes from the type-III secretion systems (**Figure 6B**). In contrast, genes encoding resistance to distinct antibiotic classes (e.g., beta-lactams, aminoglycosides, and sulphonamides) were often positively correlated in the ICEs/IMEs from both phylogroups, consistent with the previous observations that these genes tend to be co-localized in genetic structures named integrons ^66^. Virulence genes involved in flagellar motility were also often correlated, either additionally with (phylogroup B) or without (phylogroup A) genes involved in chemotaxis ^67^. Defence systems BREX and AbiEii ^62,63^ were positively correlated in ICEs/IMEs from phylogroup B. AMR and defence systems showed a high density in ICEs/IMEs from phylogroups A and B, and their frequencies were positively correlated in both phylogroups.

**Figure 6.**
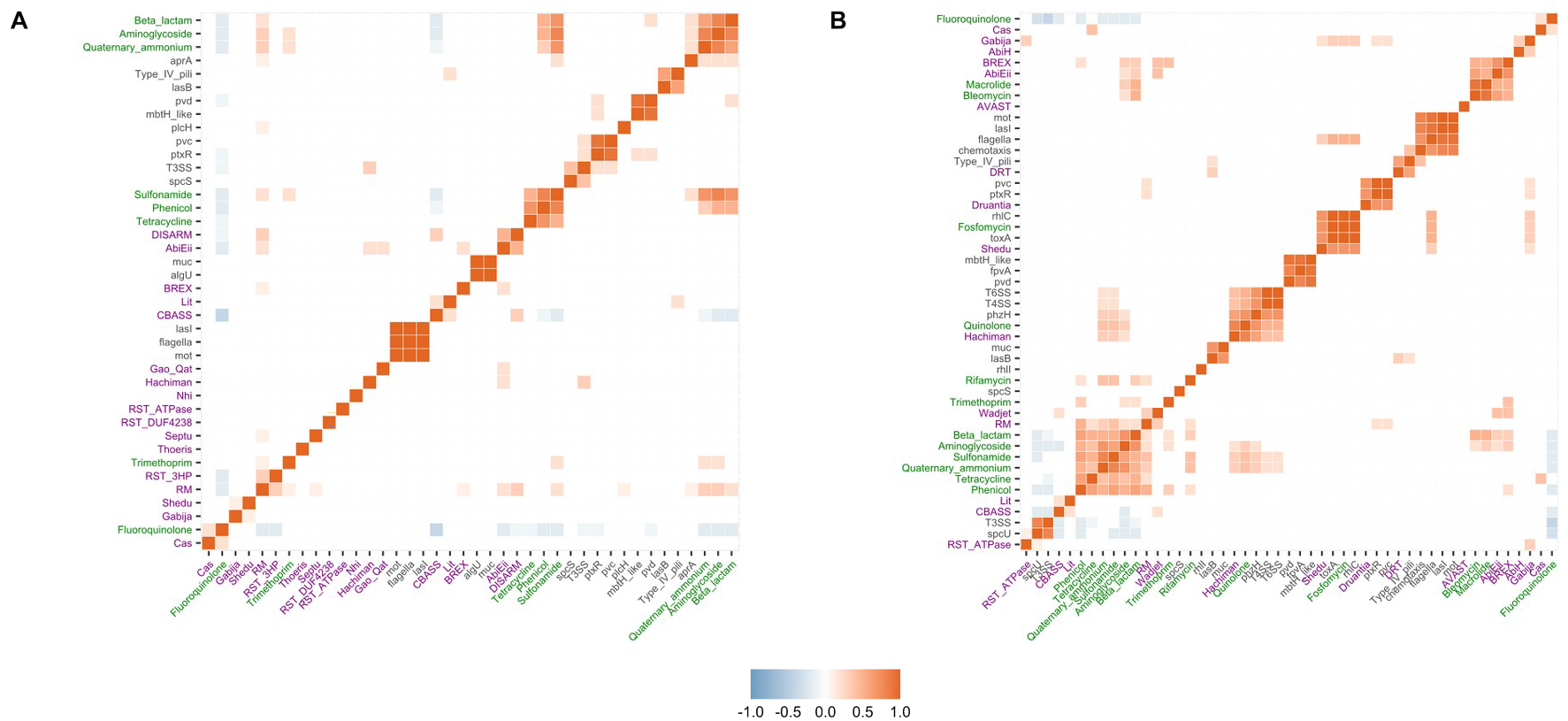
Correlation plots between AMR classes, virulence genes, and defence systems across ICEs/IMEs from phylogroup A **(A)** and phylogroup B **(B)**. The distribution of cargo genes across ICEs/IMEs was converted into a presence/absence matrix. Correlation matrices were ordered using the hierarchical clustering function. Positive correlations are shown in different shades of red, while negative correlations are shown in different shades of blue. AMR genes and point mutations encoding resistance to particular AMR classes are part of the AMRFinder database ^39^, defence systems of defense-finder ^41^, and virulence genes of the VFDB ^40^. Virulence gene labels are coloured in black, AMR in green, and defence systems in purple.

### ICEs/IMEs and RGPs from different phylogroups share high genetic similarity

We next used an alignment-free sequence similarity comparison of the ICEs/IMEs to infer an undirected network. The density plot showed a right-skewed distribution of pairwise distance similarities where the vast majority of ICE/IME pairs shared little similarity, with a Jaccard Index value below 0.5 (**Figure S15**), in accordance with the high diversity frequently observed across MGEs ^68^. To reduce the density and increase the sparsity of the network, we used the mean Jaccard Index between all pairs of RGPs as a threshold (0.12184). The network assigned 95.8% (530/553) of the ICEs/IMEs into 15 clusters (**Figure 7**). Nearly half of the ICEs/IMEs were grouped in cluster 1 (259/530, **Table S15**), which includes representatives of the three phylogroups.

**Figure 7.**
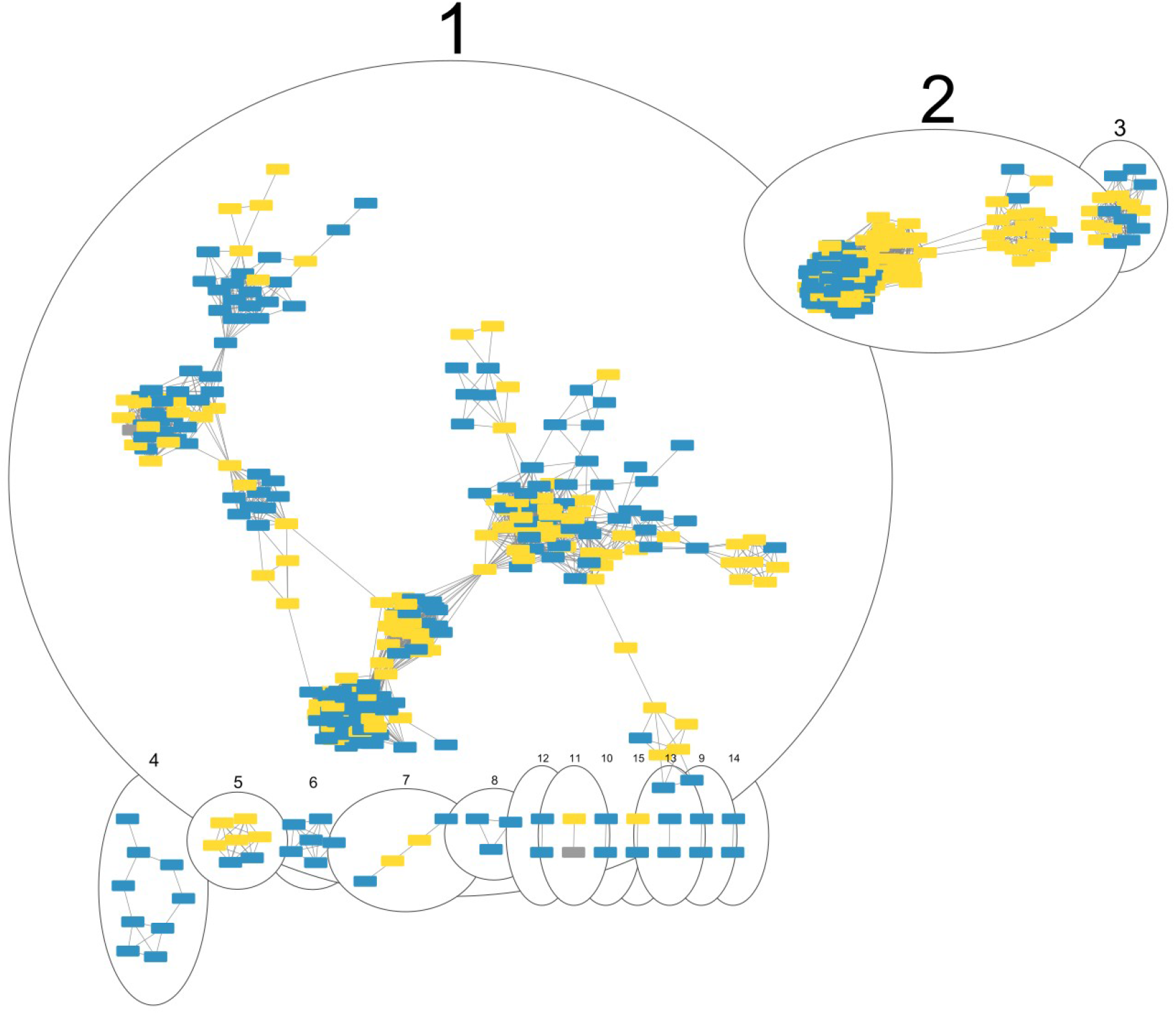
Network of clustered ICEs/IMEs from the three phylogroups, using the mean Jaccard Index between all pairs of ICEs/IMEs as a threshold. Each ICE/IME is represented by a node, connected by edges according to the pairwise distances between all ICE/IME pairs. Numbered ellipses represent ICEs/IMEs that belong to the same cluster. The network has a clustering coefficient of 0.794, a density of 0.099, a centralization of 0.217, and a heterogeneity of 0.785. ICEs/IMEs from phylogroup A are coloured in blue, from phylogroup B in yellow, and from phylogroup C in grey.

We then focused our analysis on the RGPs we extracted from all phylogroups (57901 RGPs in total). We filtered out RGPs smaller than 10kb, and calculated the Jaccard Index between all pairwise of the resulting 32744 RGPs. To reduce the density and increase the sparsity of the network, we used as a threshold the mean value (0.0919429) of the estimated pairwise distances between the 32744 RGPs identified in this study. The network assigned 99.7% (32651/32744) of the RGPs larger than 10kb into 51 clusters (**Figure S16**). While the majority of the RGP clusters were homogeneous for a given phylogroup, we also observed DNA sharing events between different phylogroups. These findings suggest that RGPs and ICEs/IMEs from different *P. aeruginosa* phylogroups share high genetic identity.

## Discussion

In this work, we explored the pangenome of the opportunistic human pathogen *P. aeruginosa* in consideration of its three main phylogroups. This approach allowed us to characterize defining properties of each phylogroup. In particular, we identified genes that are prevalent in the small phylogroup C and absent from members of the two larger phylogroups. These genes would have been classified to be part of the accessory genome in conventional analyses of the pangenome of the species as a whole. In contrast, our refined approach suggests that these genes have an evolutionary advantage in a specific genetic context that is particular to this phylogroup ^69^. Moreover, phylogroup C is also clearly distinct from the other two phylogroups A and B in having a significantly smaller genome size and a low relative frequency of AMR and defence systems across RGPs. In addition, our results indicate an inverse association between size of the phylogroup B accessory genome and presence of CRISPR-Cas systems. This association could (but need not) be causal, such that a low prevalence of CRISPR-Cas defence systems may possibly favour an increase in the size of the accessory genome. Remarkably, genomes devoid of CRISPR-Cas systems in phylogroups A and B were generally significantly larger than those with these systems, a trend that was no longer observed when only considering the non-RGP (“masked”) genomes. This observation is consistent with the hypothesis that CRISPR-Cas systems can constrain horizontal gene transfer in *P. aeruginosa* ^50–52,70^, at least for genomes belonging to the larger phylogroups.

The three phylogroups vary substantially in the distribution of AMR genes, defence systems, and virulence genes. This variation is particularly apparent in the separate analyses of RGPs and masked genomes. While the length of RGPs is substantially smaller than that of masked genomes, the absolute counts of most defence systems were higher in RGPs than in masked genomes across the three phylogroups (**Figure S11**). Curiously, representatives of the recently described set of defence systems that are part of Doron’s seminal study ^61^, such as Zorya, Wadjet, and Hachiman systems, were exclusively found in RGPs across the three phylogroups. In Doron’s study, the authors demonstrated that the Wadjet system provided protection against plasmid transformation in *Bacillus subtilis,* while the Zorya and Hachiman systems mediated defence against bacteriophages. These findings highlight the important role of defence systems encoded in RGPs in protecting genomes against infection by foreign DNA and their contribution to MGE-MGE conflict. Moreover, AMR and defence systems are rare in RGPs from phylogroup C, which may suggest that these strains are more often subjected to infection by foreign DNA. Assuming that there is no sampling bias across the three phylogroups, then the smaller number of phylogroup C members in public databases could thus be a consequence of the weaker arsenal of AMR and defence systems. Alternatively, phylogroup C strains may indeed be underrepresented, for example if they mainly occur in non-clinical habitats, which are usually less well sampled. Collecting *P. aeruginosa* samples from distinct geographic regions and environments may further help us reconstruct variation in metabolic competences and their connection to origin ^71^.

In general, our results underscore the role of ICEs/IMEs as vectors not only of AMR genes ^59^, but also of defence systems. Indeed, most of these systems show nonrandom clustering in defence islands and are often co-localized with mobilome genes ^61,72–74^. Co-occurrence of genes alone, however, does not infer an ecological interaction between them ^75^. Recently, it was proposed that the accessory genome of the genus *Pseudomonas* is influenced by natural selection, showing a higher level of genetic structure than would be expected if neutral processes governed the pangenome formation ^76^. This suggests that coincident genes in ICEs/IMEs are more likely to act together for the benefit of the host or to ensure their own maintenance ^9,11^. ICEs/IMEs, in particular, provide abundant material for the experimental study of bacterial defence systems. For example, SXT ICEs in *Vibrio cholerae,* which are also involved in AMR, consistently encode defence systems localized to a single hotspot of genetic shuffling ^77^. Additionally, ICEs in *Acidithiobacillia* carry type-IV CRISPR-Cas systems with remarkable evolutionary plasticity, which are often involved in MGE-MGE warfare ^78^. Moreover, a recent study proposed that size constraints may account for the low abundance of large defence systems on prophages ^41^. In turn, the comparatively larger size of ICEs/IMEs (when compared with prophages) ^79^ may then explain that they commonly harbor large systems such as BREX and defence island system associated with restriction–modification (DISARM) ^80^ across our dataset (**Figure S13**). Even though the CBASS systems are not as prevalent as restrictionmodification and CRISPR-Cas systems across the bacterial phylogeny ^41^, three types of this system were found across ICEs/IMEs from the larger phylogroups.

For our analyses, we used complete and draft genome assemblies retrieved from public databases. However, incomplete genome assemblies likely impact RGP definition, due to highly fragmented genomes, that might have inadvertently split RGPs into multiple contigs. With that in mind, we subsampled the complete genomes from our collection and used these to accurately delineate ICEs/IMEs. With the sequence similarity comparison between all pairs of ICEs/IMEs found in this study, as well as between all pairs of RGPs, we were able to explore interactions between these elements, suggesting that members of the same and of different phylogroups frequently undergo DNA shuffling events. Importantly, this network-based approach using pairwise genetic distances of alignment-free *k*-mer sequences between MGE pairs has bypassed the exclusion of non-coding elements, providing a more comprehensive picture of MGE populations and dynamics ^51,81^. Nevertheless, with the current progress in sequence technology, especially including long-read sequencing, we envision a much larger number of completely assembled *P. aeruginosa* genomes in the future, which will then improve reliable assessment of the RGP composition and the role of particular MGEs or gene functions in shaping this species’ genome characteristics.

To conclude, our work used a refined approach to explore phylogroup-specific and pangenome dynamics in *P. aeruginosa*. Members of phylogroup B contribute a comparatively larger number of pangenome families, have larger genomes, and have a lower prevalence of CRISPR-Cas systems. AMR and defence systems are pervasive in RGPs and ICEs/IMEs from phylogroups A and B, and these two functional groups are often significantly correlated, including both positive and negative correlations. We also observed multiple interaction events between the accessory genome content both between and within phylogroups, suggesting that recombination events are frequent. Our conclusions are contingent on the current range of sequenced genomes for *P. aeruginosa.* We cannot exclude that some groups, for example phylogroup C and possibly its subgroups, are not fully represented in the currently available data. Future sequencing efforts are likely to rectify such a possible problem, thus allowing to test the findings from our study. Finally, our work provides a representative set of phylogenetically diverse *P. aeruginosa* strains, the mPact strain panel, which should prove useful as a reference set for future functional analyses. Such functional analyses may help to experimentally assess the underlying reasons for some of the correlations identified in our study, for example the role of specific defence systems in RGP size expansion or in mediating conflict between different MGE types.

## Supporting information

Captions for supplementary tables

## Declaration of interests

We declare no competing interests.

## Contributors

JB conceptualized and designed the work, acquired and analysed the data, interpreted the data, and wrote the original draft of the manuscript. LT conceptualized and designed the work, acquired the data, and interpreted the data. JF, FB, CU, JK, SN, and BT acquired and analysed the data. HS conceptualized and designed the work, interpreted the data, and contributed to writing of the original draft of the manuscript. HS, SN, and BT acquired funding for this work. All authors read, revised, and approved the final manuscript.

## Data sharing

Scripts for reproducing the analyses performed in this work are available at https://gitlab.gwdg.de/botelho/pa_pangenome. The representative set of *P. aeruginosa* genomes and the input file used for the network analysis in Figure 7 are available at the Figshare project https://figshare.com/projects/P_aeruginosa_pangenome/155021. Analyses were made with a combination of shell and R 4.0.3 scripting. Sequencing performed in this project were deposited in NCBI under the Bioproject accession number PRJNA810040.

## Acknowledgments

We acknowledge financial support from the German Science Foundation (grant SCHU 1415/12-2 to HS, funding under Germany’s Excellence Strategy EXC 2167-390884018 as well as the Research Training Group 2501 TransEvo to HS and SN, and funding within the SFB 900 TP A2 to BT), the Max-Planck Society (Max-Planck fellowship to HS), the Leibniz ScienceCampus Evolutionary Medicine of the Lung (EvoLUNG, to HS and SN), and the BMBF program Medical Infection Genomics (AZ 0315827A to BT). This work was supported by the Kiel Life Science Postdoc Award to JB and by the DFG Research Infrastructure NGS_CC (project 407495230) as part of the Next Generation Sequencing Competence Network (project 423957469). NGS was carried out at the Competence Centre for Genomic Analysis (Kiel). This research was supported in part through high-performance computing resources available at the Kiel University Computing Centre.

## SUPPLEMENTARY FIGURES

**Figure S1.**
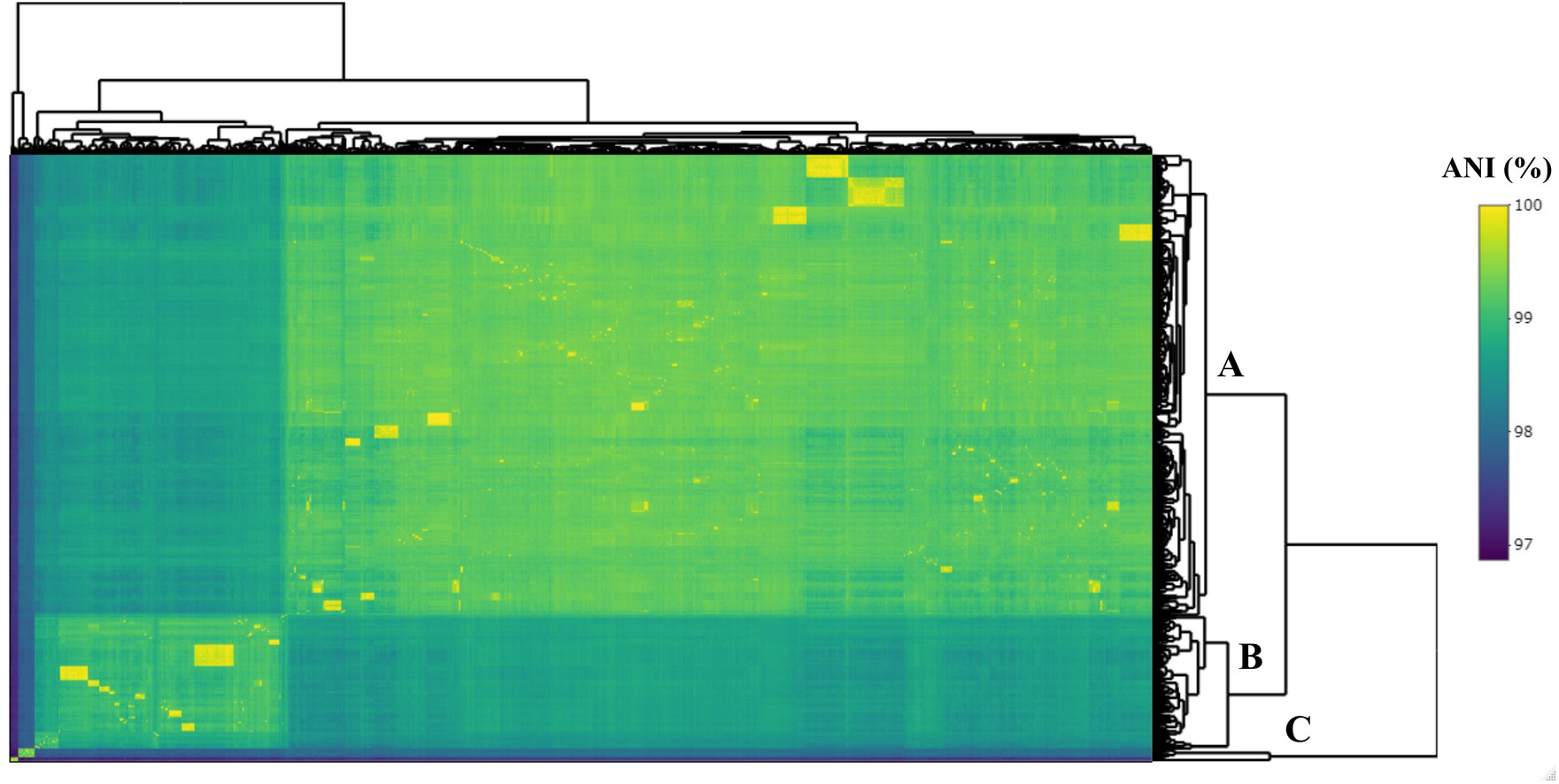
Matrix of Average Nucleotide Identity (ANI) scores between the 2009 *P. aeruginosa* genomes used in this study. Row and column dendrograms are displayed. Hierarchical clustering was performed with the complete-linkage clustering method. The three phylogroups identified in this study are highlighted in the row dendrogram.

**Figure S2.**
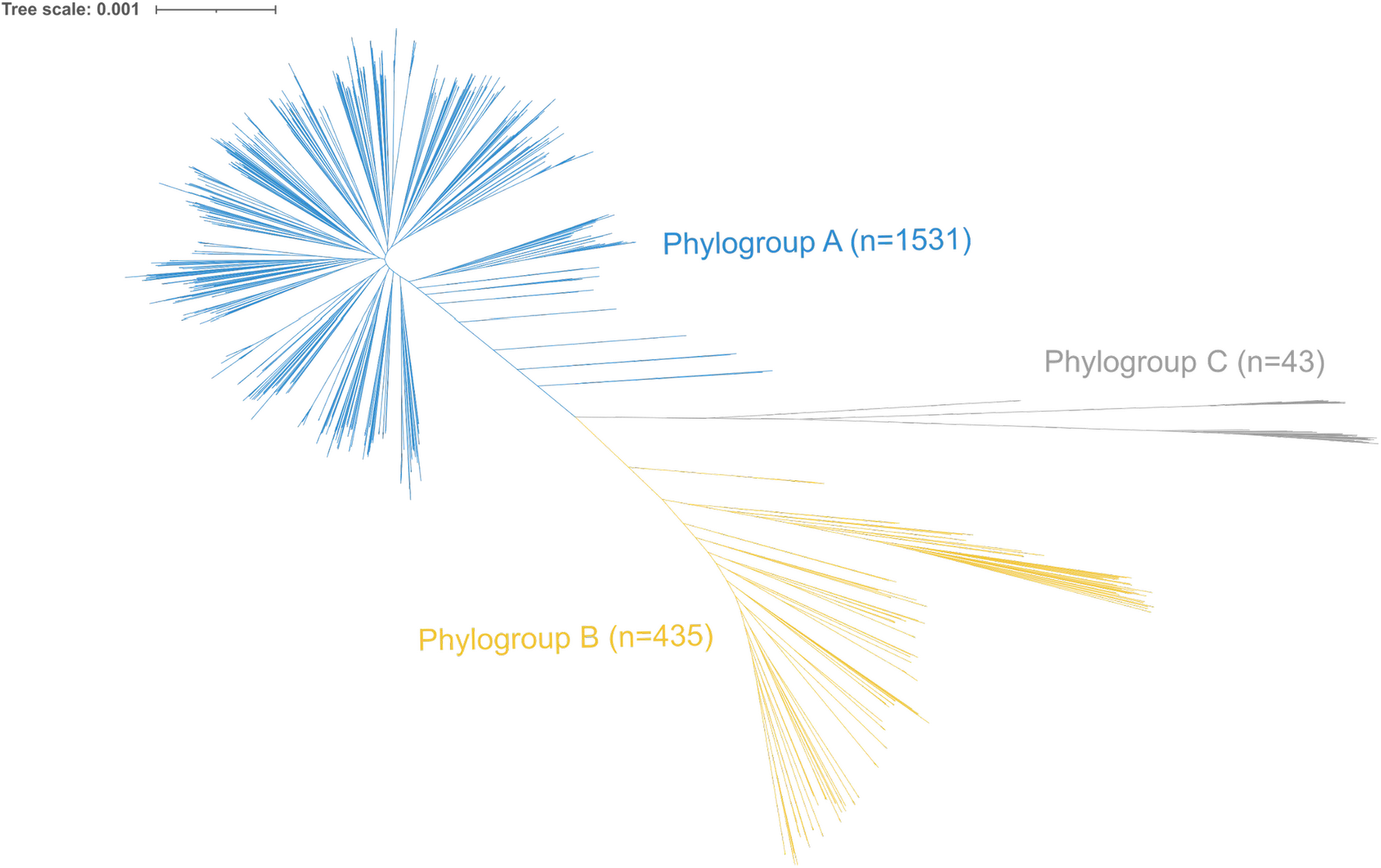
Maximum-likelihood tree of the softcore-genome alignment of all *P. aeruginosa* isolates used in this study (n=2009), corrected for recombination. The scale bar represents the genetic distance. Members of phylogroup A are coloured in blue, B in yellow, and C in grey.

**Figure S3.**
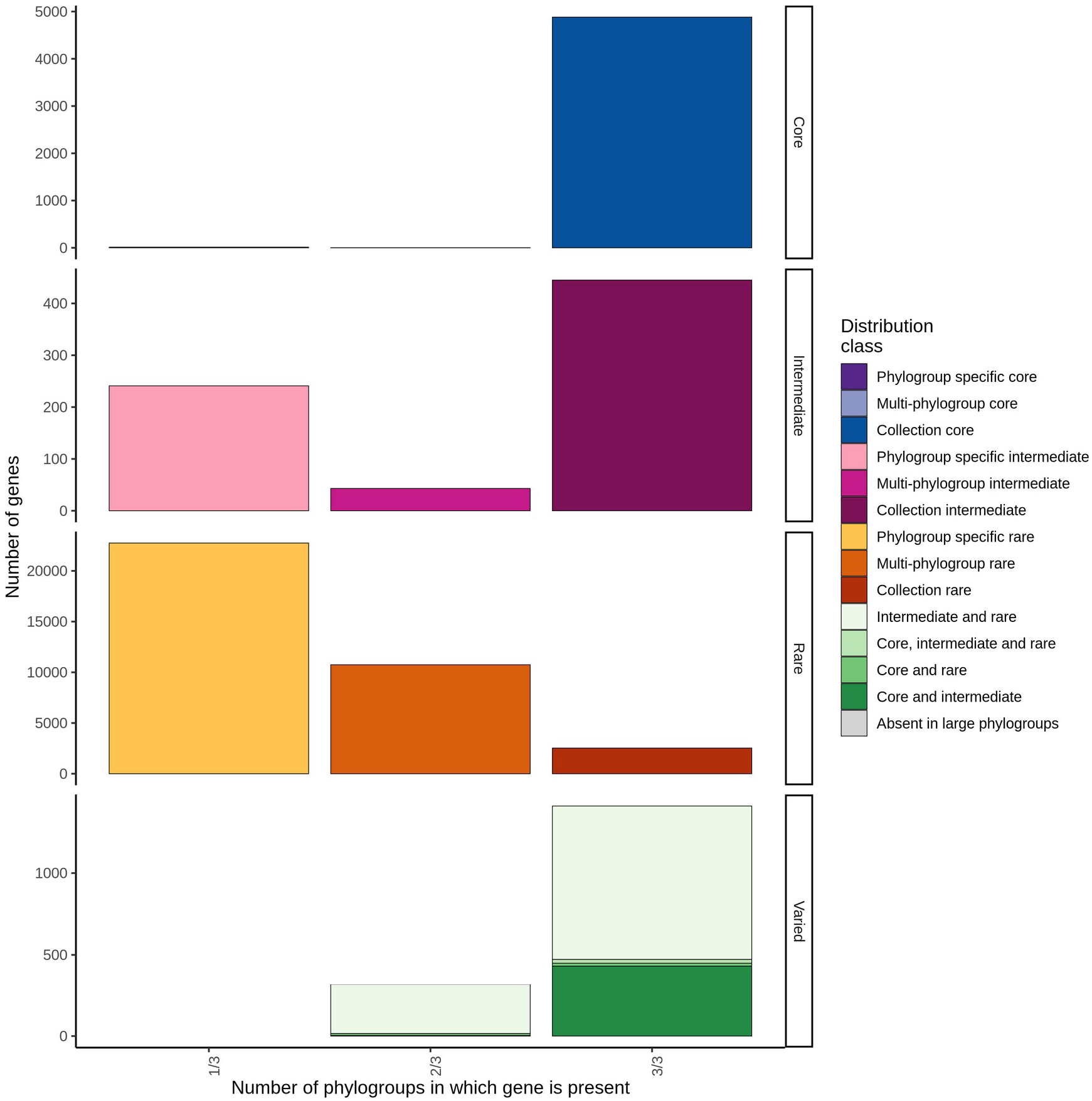
Barplots of the distribution of gene families into core, intermediate, rare, or varied parts of the pangenome across phylogroups. The first column shows genes that are specific to a given phylogroup, and further classified into core (≥95%), intermediate, rare (≤15%), or varied. The second column shows genes that are specific to two phylogroups, and their classification into core, intermediate, rare, or varied. The third column shows genes that are present across all three phylogroups, and their classification into core, intermediate, rare, or varied. A different colour is assigned to each classification. To create the plot, we modified the R script available in https://github.com/ghoresh11/twilight/blob/master/classify_genes.R.

**Figure S4.**
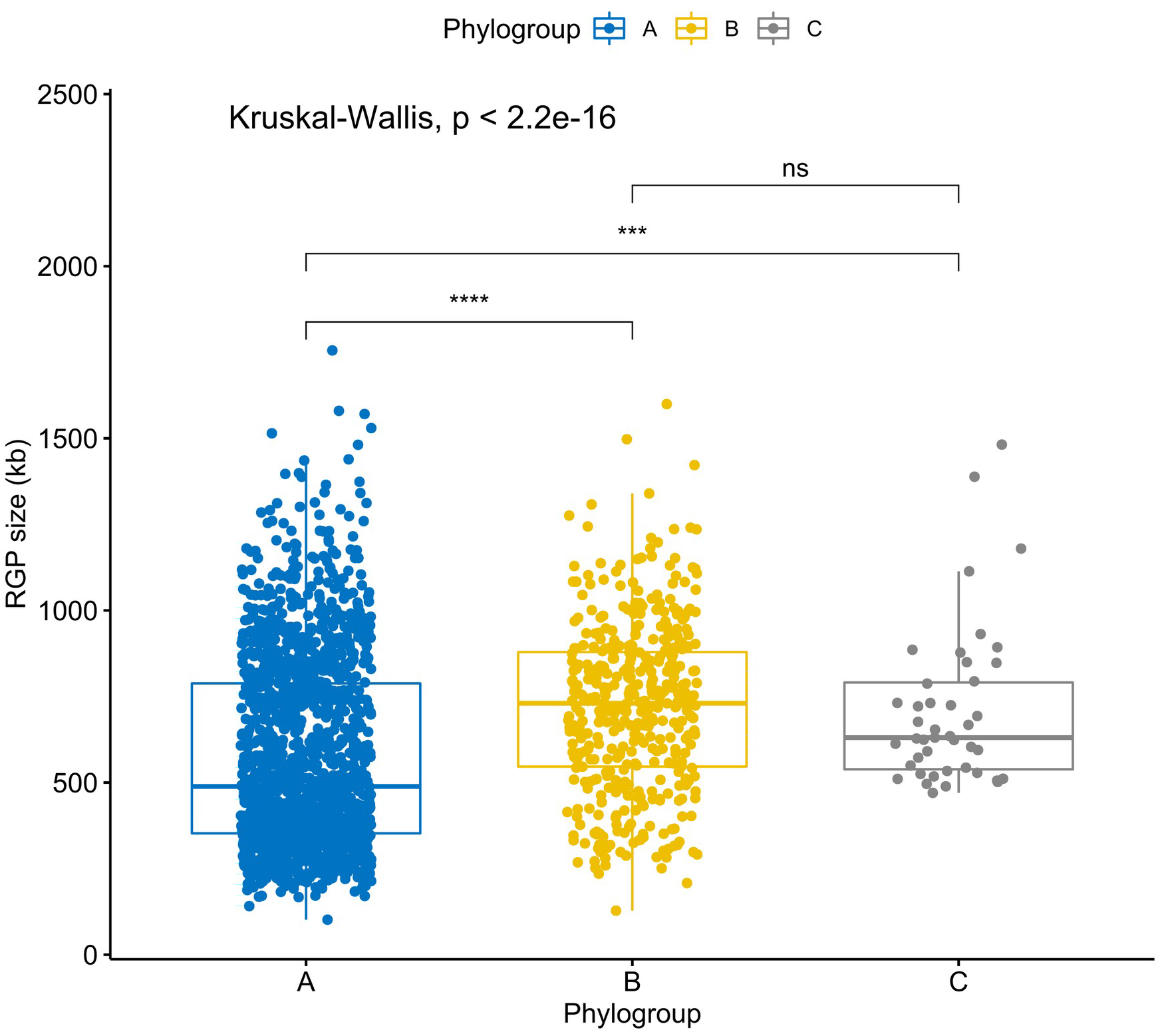
Boxplots showing the variation in RGP size across the three phylogroups. Values above 0.05 were considered as non-significant (ns). Stars indicate significance level: * p <= 0.05, ** p <= 0.01, *** p <= 0.001, and **** p <= 0.0001. Boxplots in blue represent phylogroup A, yellow B, and grey C.

**Figure S5.**
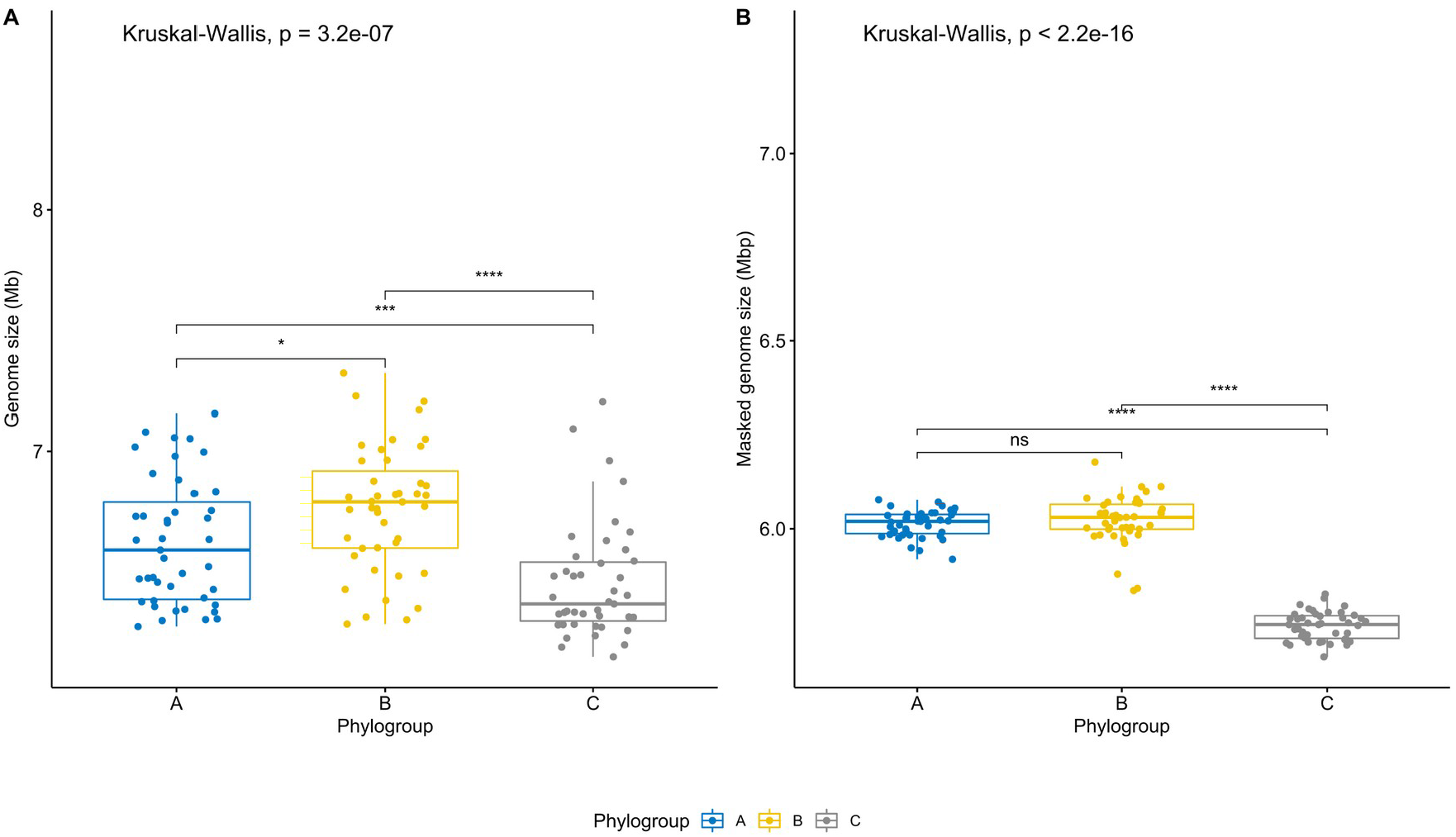
Boxplots representing the variation in genome size **(A)** and masked genome size **(B)** across the three phylogroups. Phylogroup sample sizes were adjusted to sample size of the smallest group, phylogroup C (with 43 genomes). Values above 0.05 were considered as non-significant (ns). Stars indicate significance level: * p <= 0.05, ** p <=0.01, *** p <=0.001, and **** p <= 0.0001. Boxplots in blue represent phylogroup A, yellow B, and grey C.

**Figure S6.**
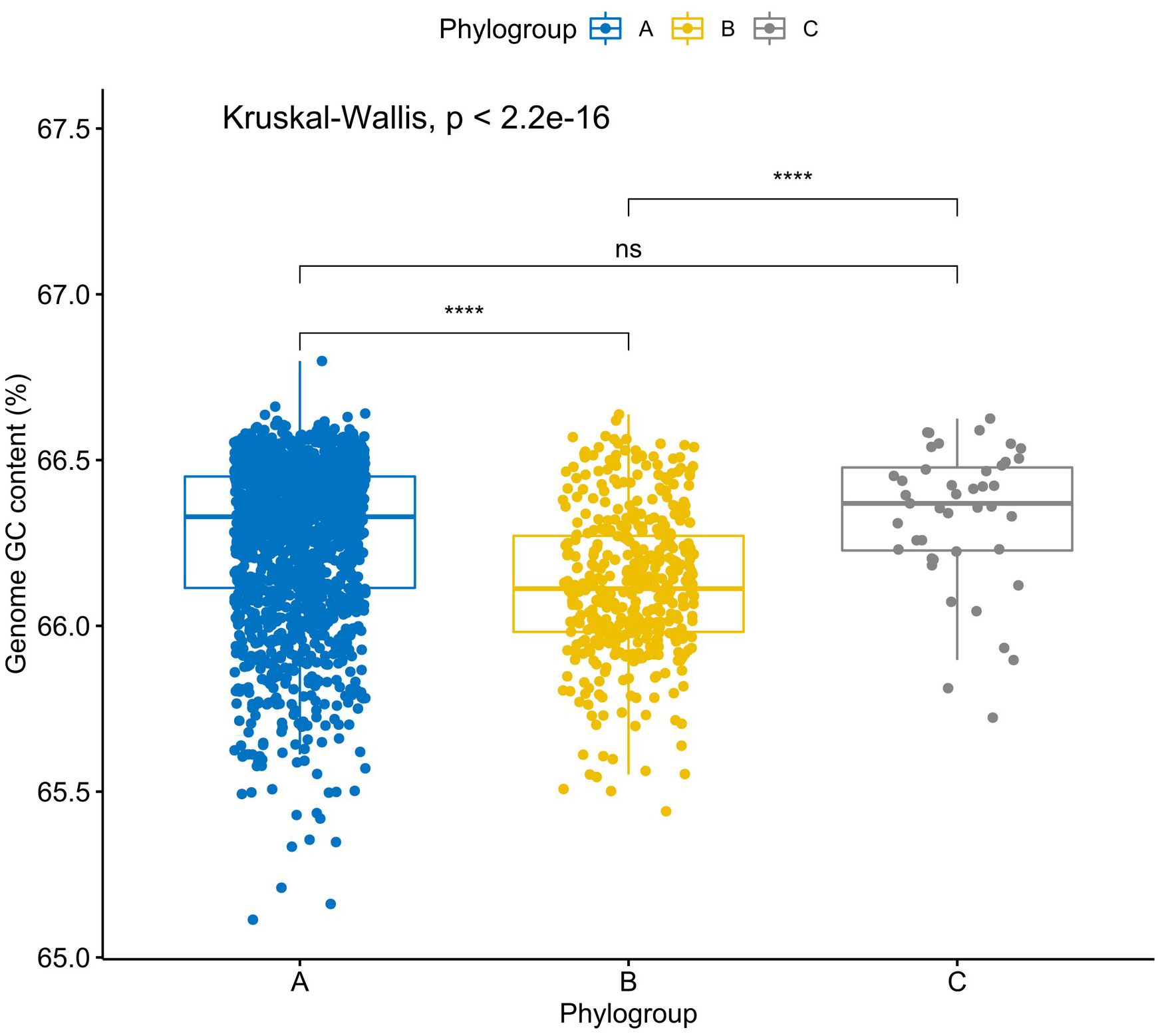
Boxplots showing the variation in GC content across the three phylogroups. Values above 0.05 were considered as non-significant (ns). Stars indicate significance level: * p <= 0.05, ** p <= 0.01, *** p <= 0.001, and **** p <= 0.0001. Boxplots in blue represent phylogroup A, yellow B, and grey C.

**Figure S7.**
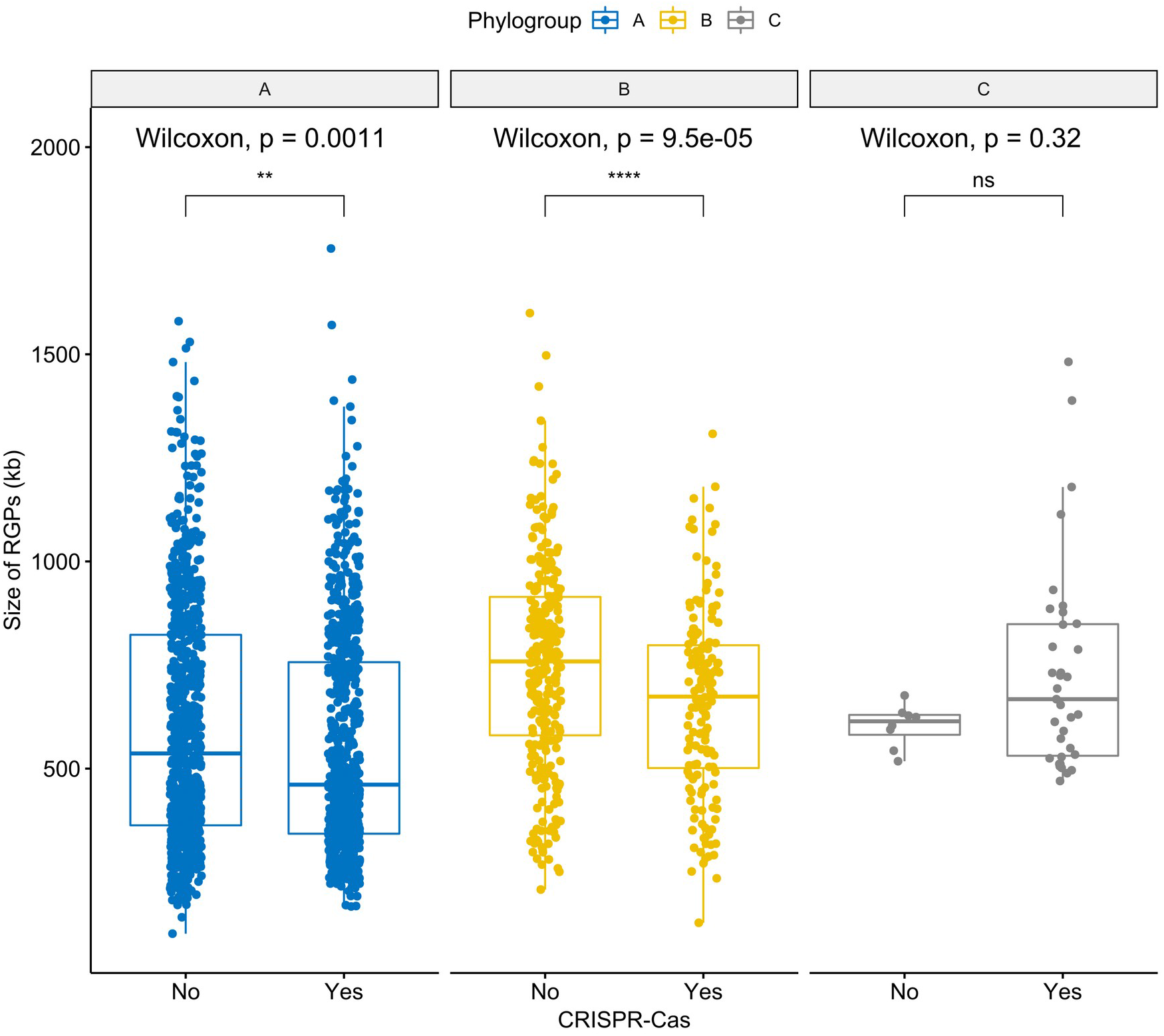
Boxplots representing the variation in the size of RGPs across pairs of conspecific genomes from the same phylogroup with and without CRISPR-Cas systems. Values above 0.05 were considered as non-significant (ns). Stars indicate significance level: * p <= 0.05, ** p <= 0.01, *** p <= 0.001, and **** p <= 0.0001. Boxplots in blue represent phylogroup A, yellow B, and grey C.

**Figure S8.**
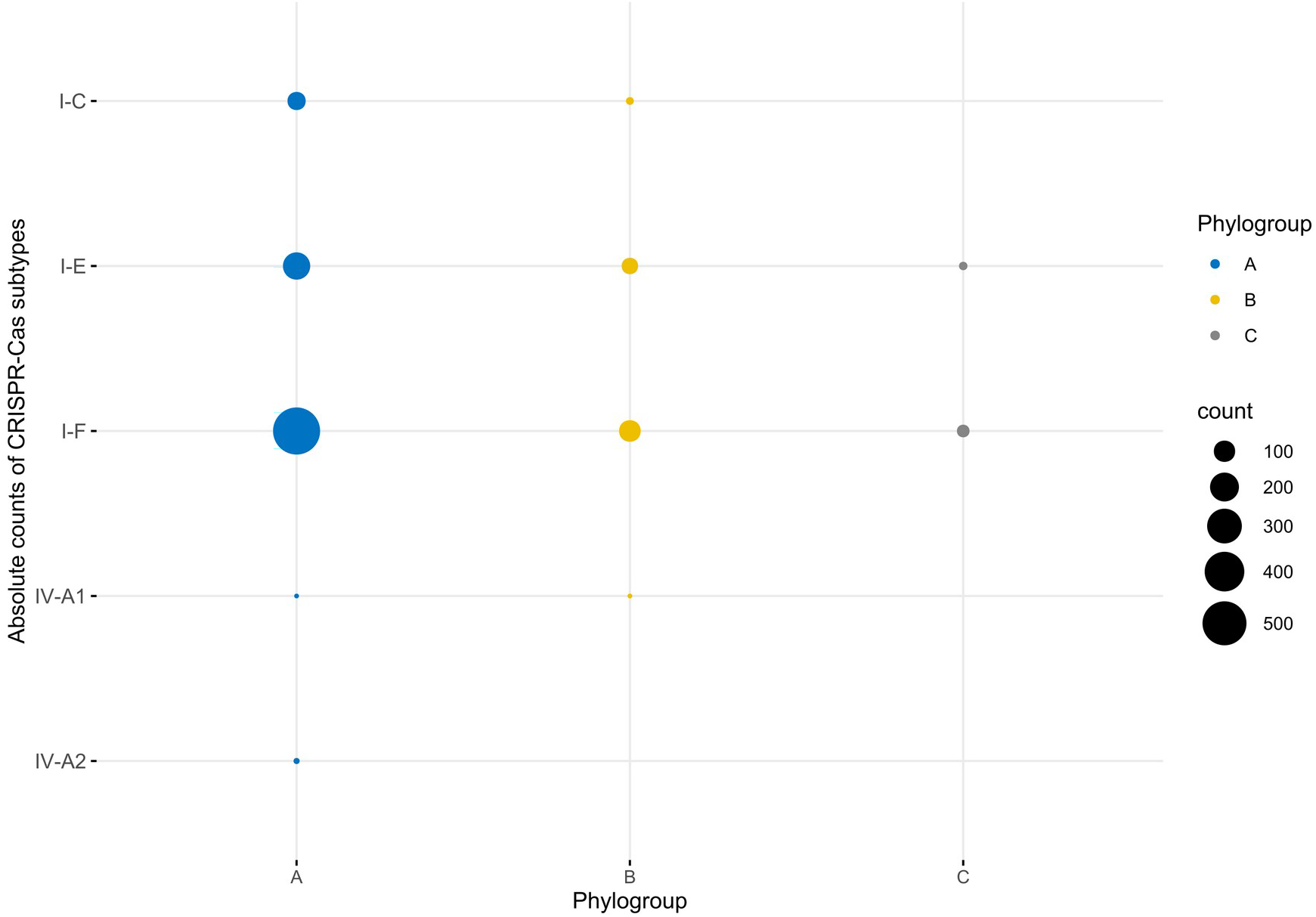
Absolute counts of CRISPR-Cas subtypes identified across genomes from the three phylogroups. Circles in blue represent phylogroup A, yellow B, and grey C. Circle size is proportional to the number of absolute counts.

**Figure S9.**
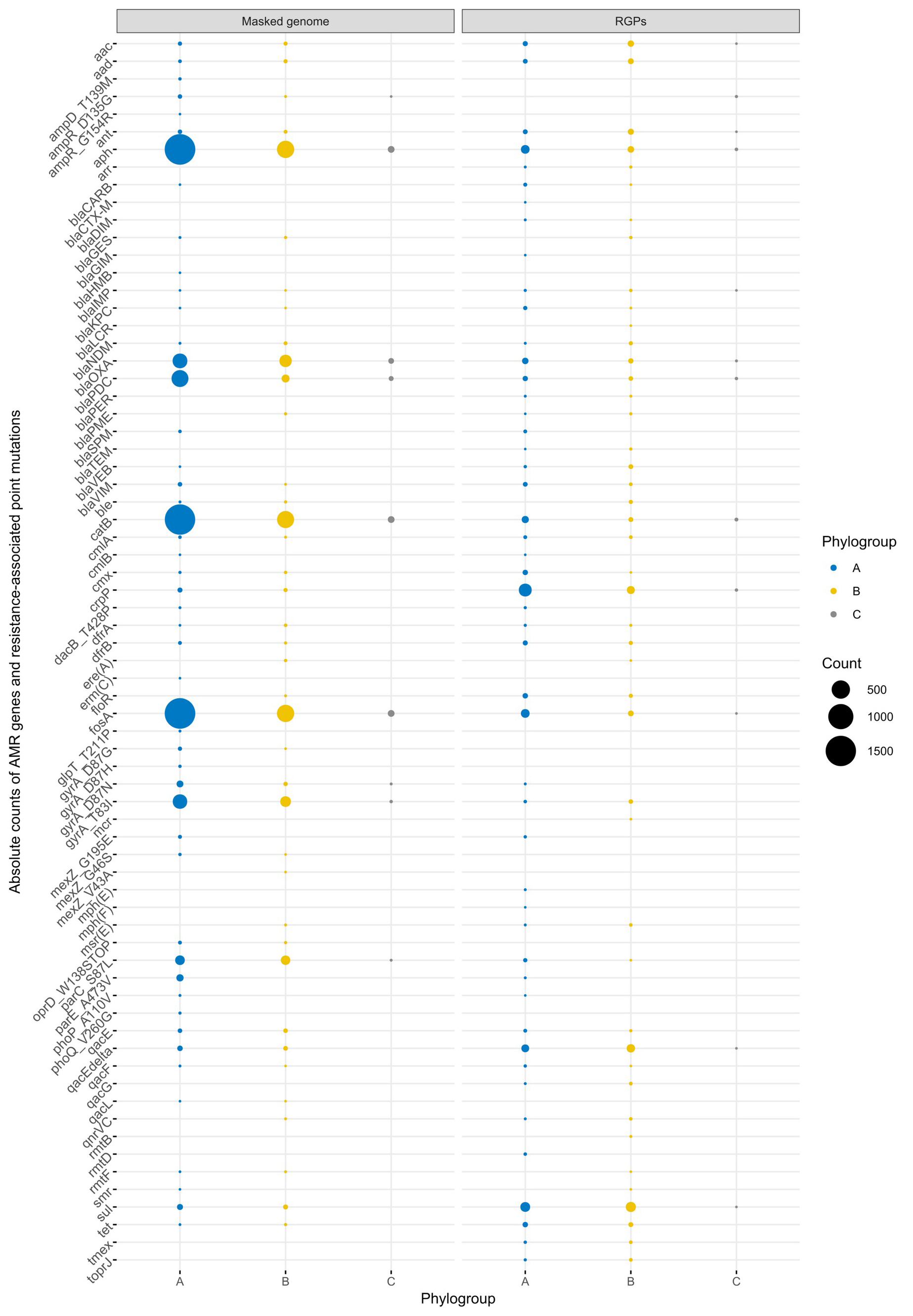
Absolute counts of AMR genes and resistance-associated point mutations across masked genomes and RGPs from the three phylogroups. Genes and mutations are part of the AMRFinder database (45). Circle size is proportional to the number of absolute counts. Circles in blue represent phylogroup A, yellow B, and grey C.

**Figure S10.**
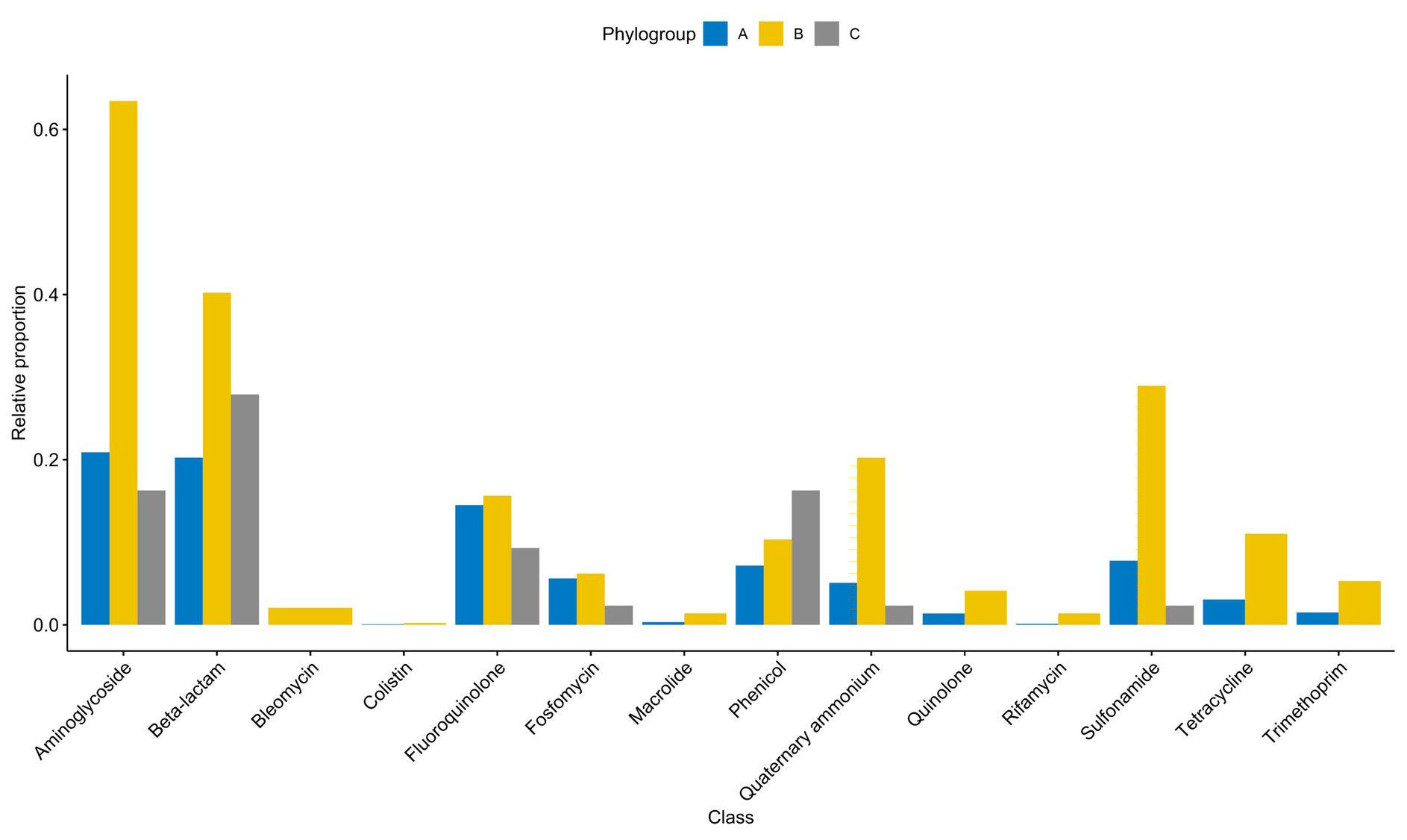
Barplots showing the relative proportion of genes encoding resistance to antibiotics from different classes across RGPs from the three phylogroups. Genes were normalized to the total number of genomes found in each phylogroup. Bars in blue represent phylogroup A, yellow B, and grey C.

**Figure S11.**
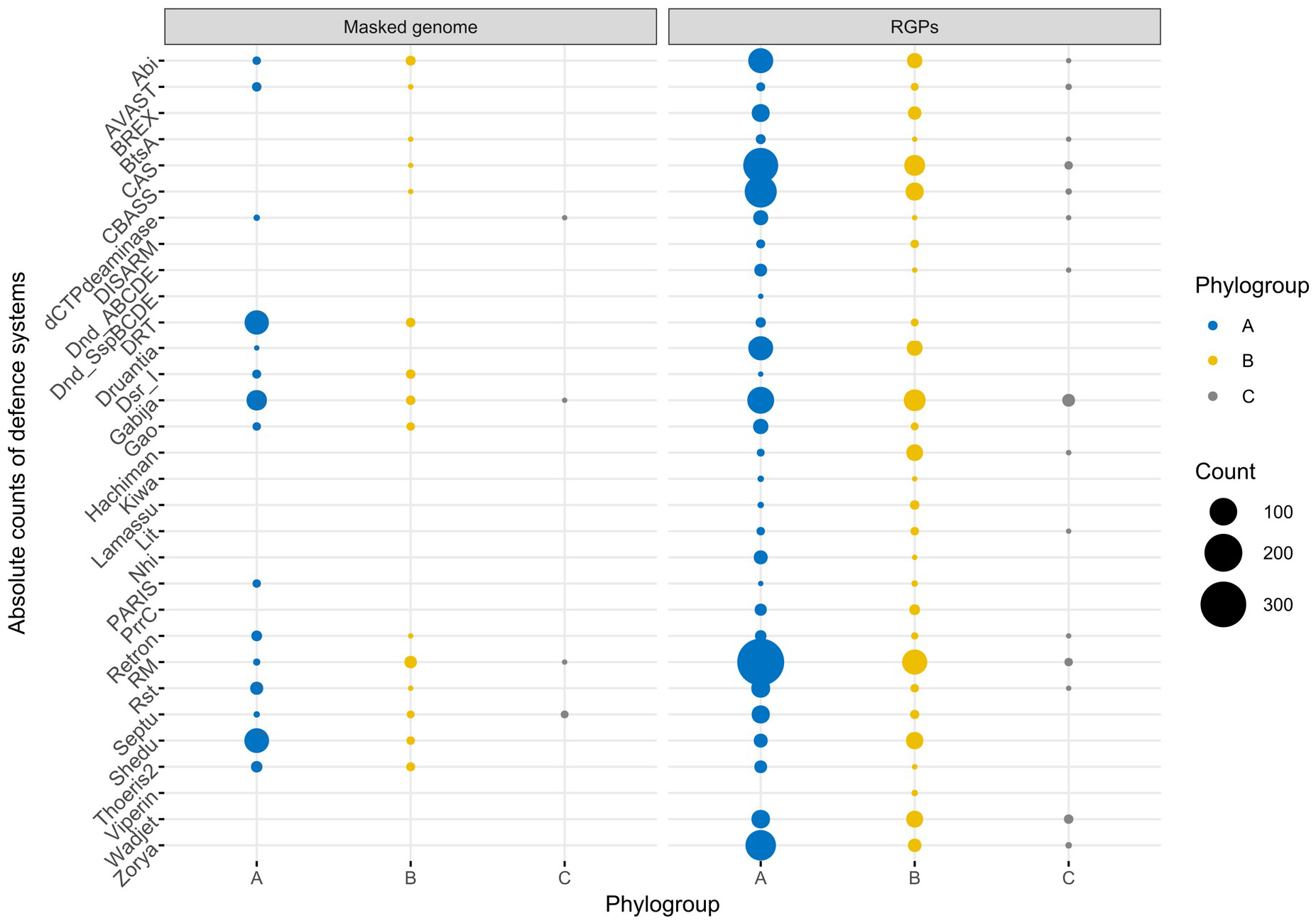
Absolute counts of defence systems across masked genomes and RGPs from the three phylogroups. Defence systems are part of the defense-finder database (36). Circle size is proportional to the number of absolute counts. Circles in blue represent phylogroup A, yellow B, and grey C.

**Figure 12.**
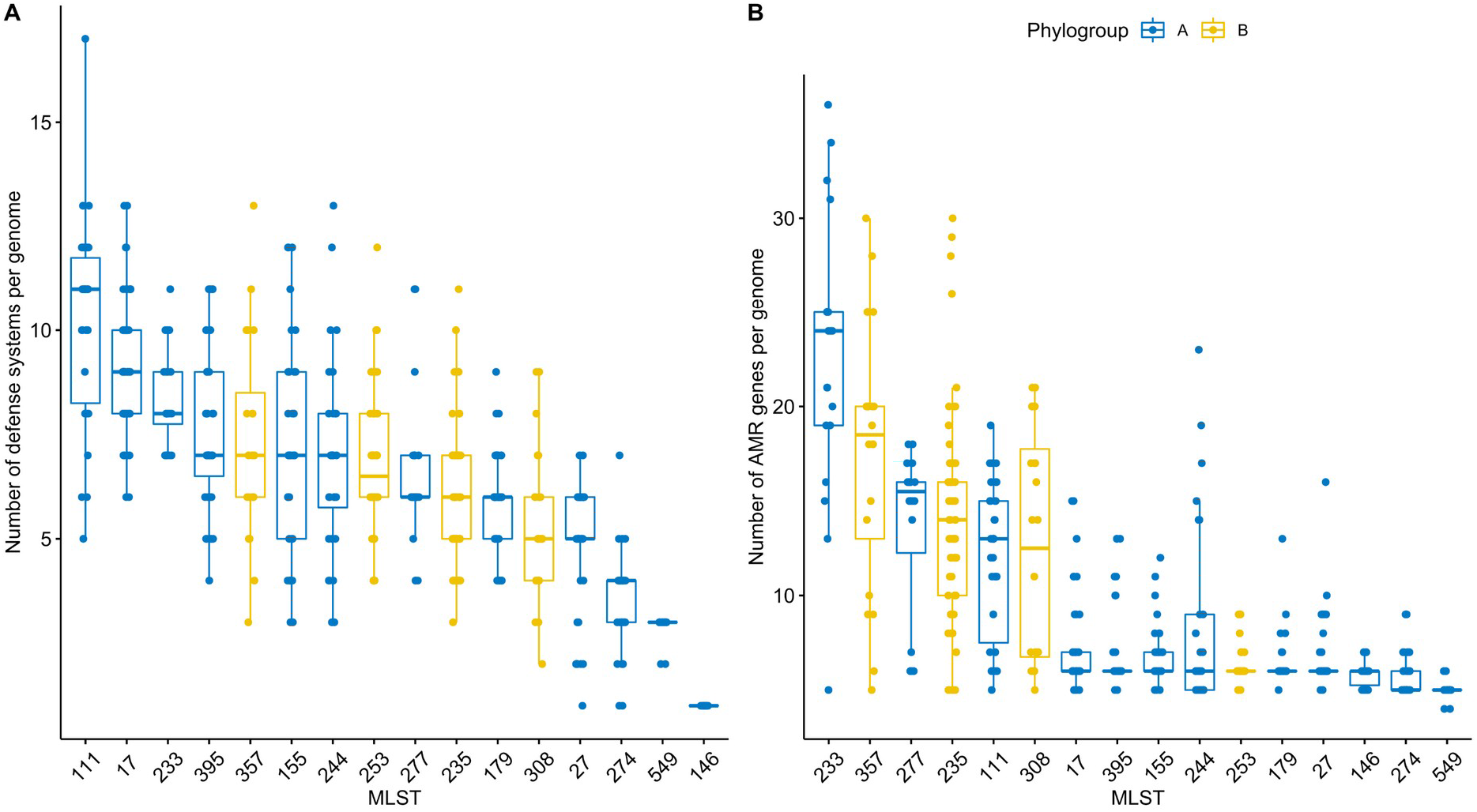
Boxplots representing the variation in the number of defense systems **(A)** and AMR genes **(B)** found across the genomes from the main MLST profiles found in this study. Only MLST profiles with at least 10 genomes are shown. Boxplots are ordered in descending order by the median values.

**Figure S13.**
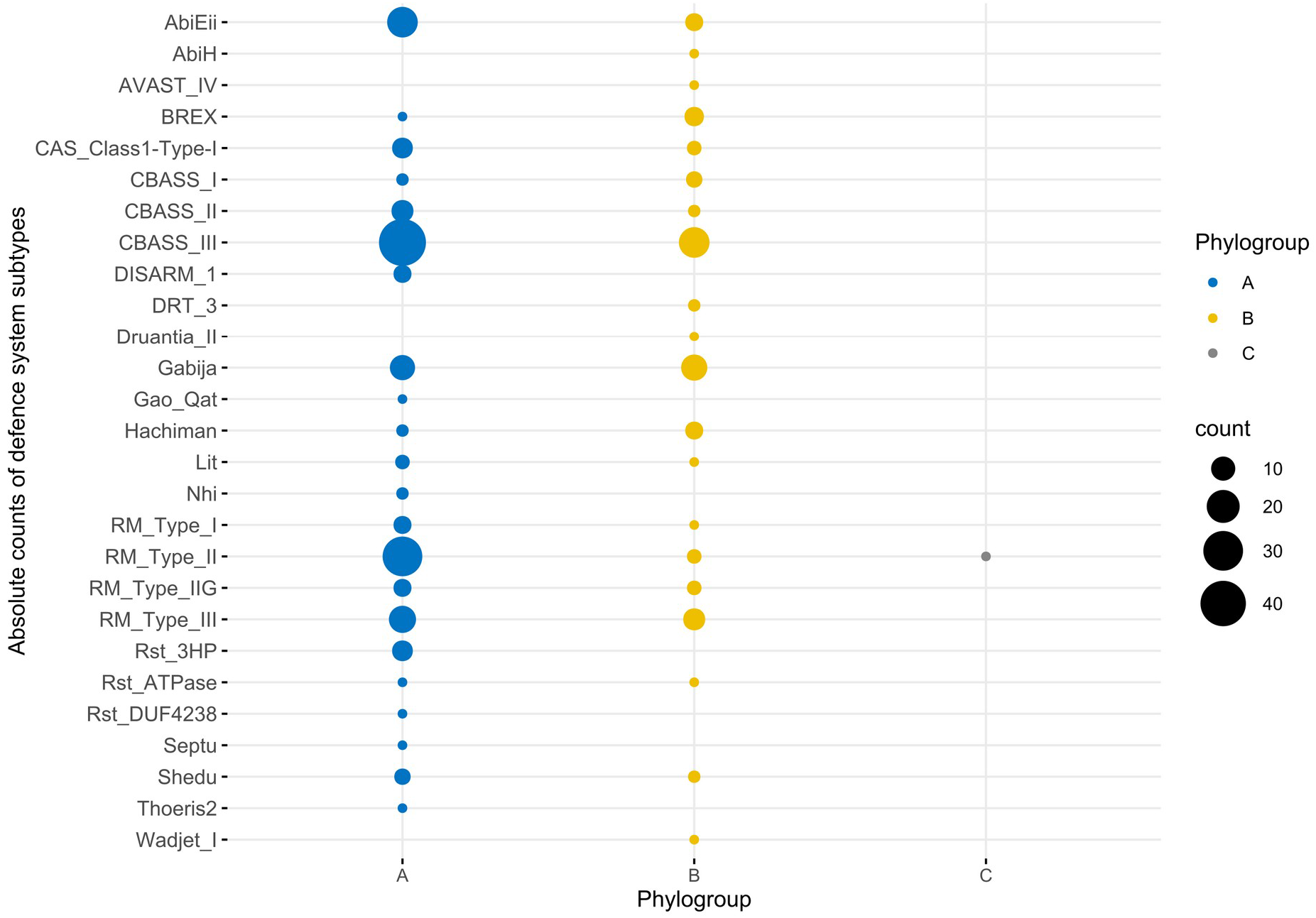
Absolute counts of defence systems across ICEs/IMEs from the three phylogroups. Defence systems are part of the defense-finder database (36). Circle size is proportional to the number of absolute counts. Circles in blue represent phylogroup A, yellow B, and grey C.

**Figure S14.**
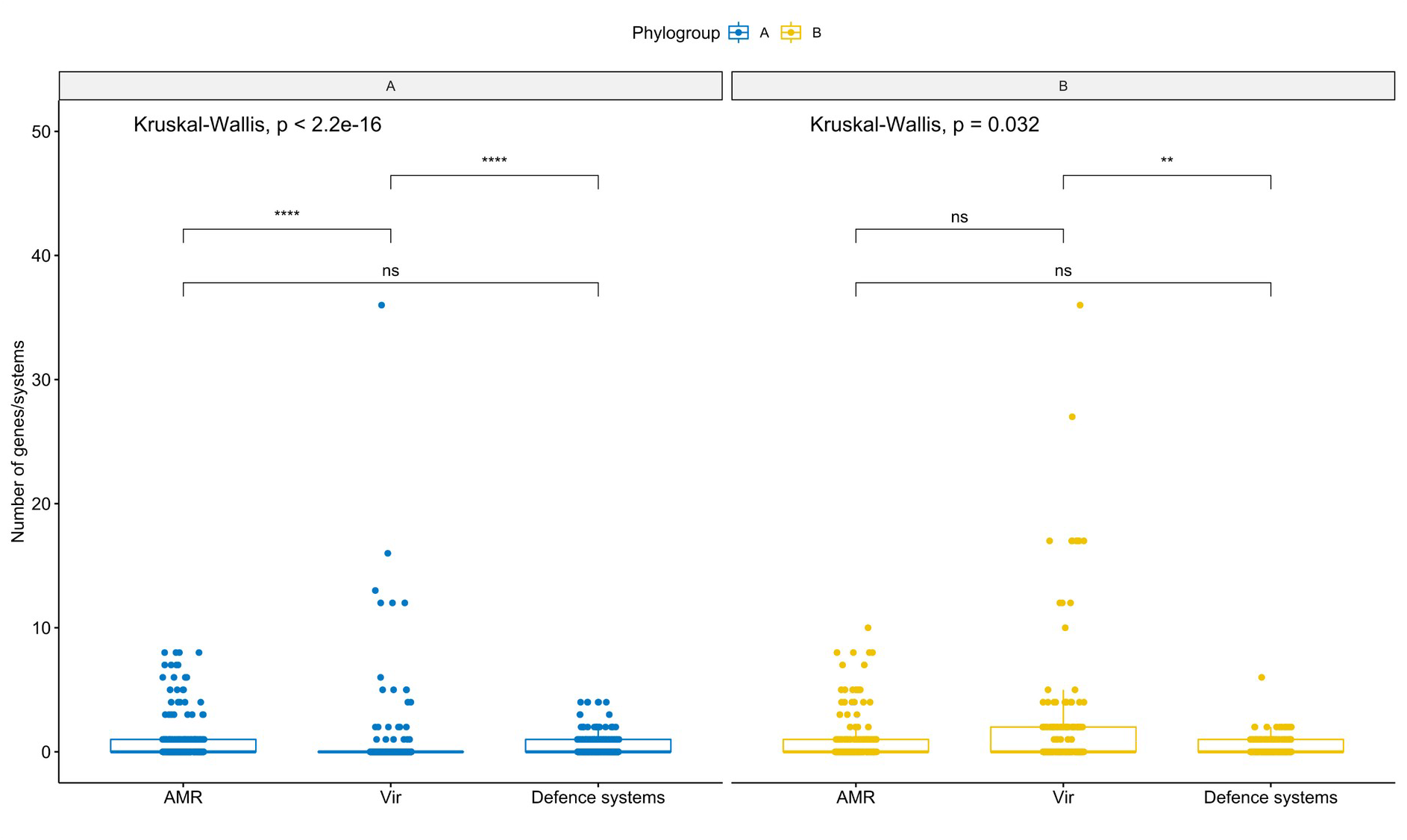
Boxplots representing the variation in the number of AMR genes, defence systems, and virulence genes found in ICEs/IMEs across the two larger phylogroups A and B. Values above 0.05 were considered as non-significant (ns). Stars indicate significance level: * p <= 0.05, ** p <= 0.01, *** p <= 0.001, and **** p <= 0.0001. Boxplots in blue represent phylogroup A, yellow B, and grey C.

**Figure S15.**
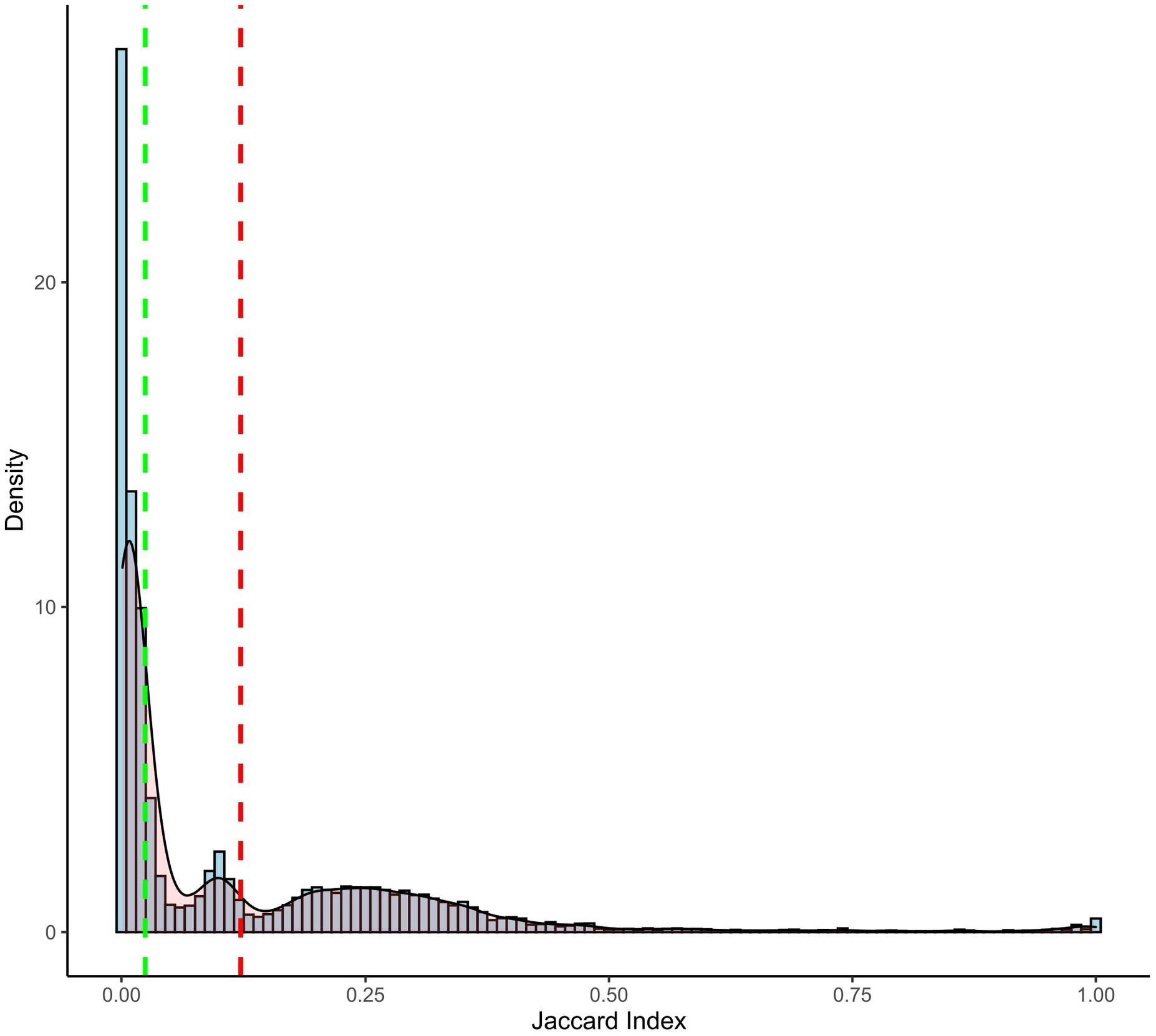
Histogram showing the right-skewed distribution of the Jaccard Index between all pairs of ICEs/IMEs. Median and mean values are highlighted by vertical dashed lines in green and red, respectively.

**Figure S16.**
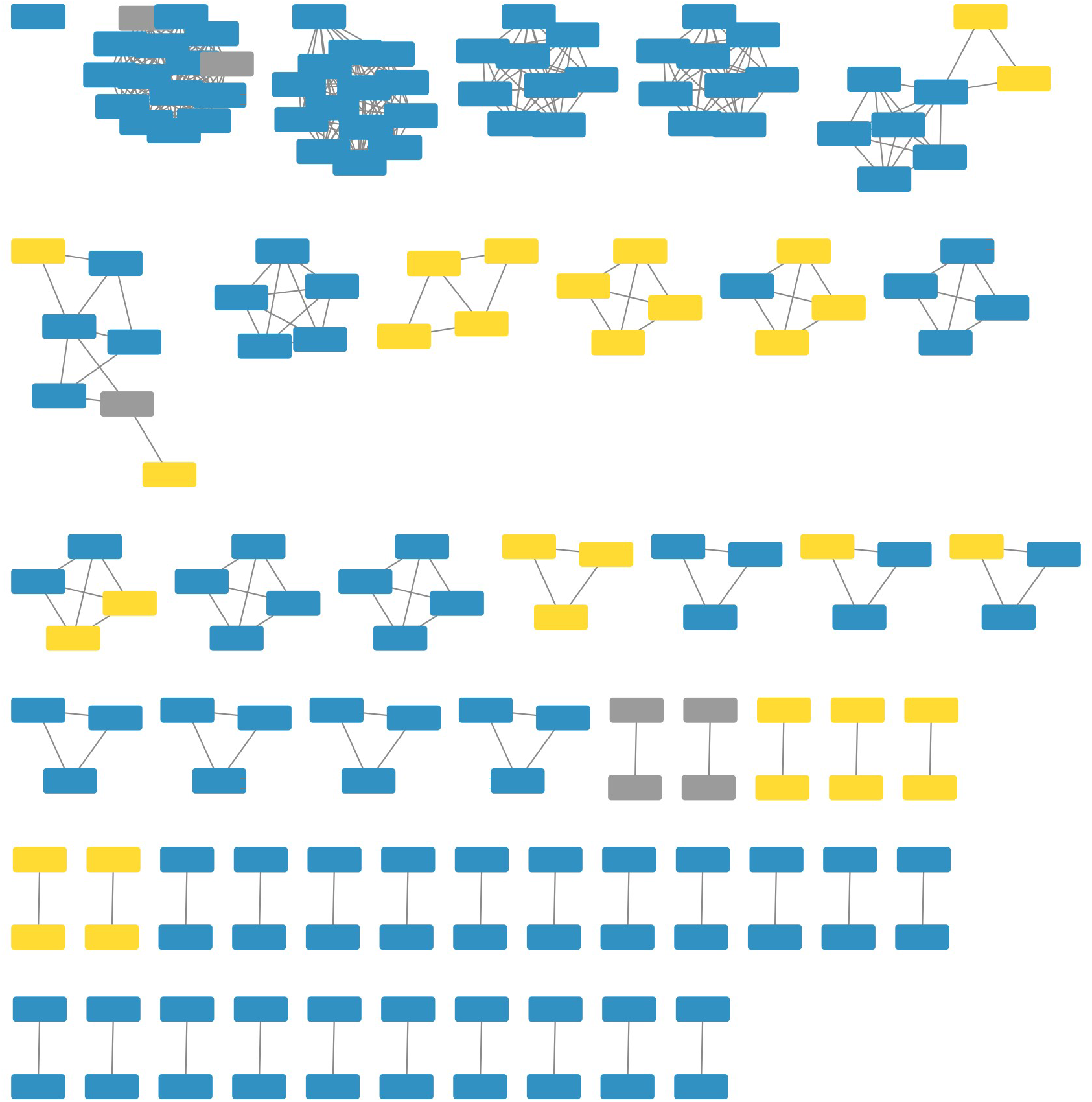
Network of clustered RGPs from the three phylogroups, using the mean Jaccard Index between all pairs of RGPs as a threshold. Each RGP is represented by a node, connected by green edges according to the pairwise distances between all RGPs pairs. Numbered ellipses represent RGPs that belong to the same cluster. The network has a clustering coefficient of 0.777, a density of 0.007, a centralization of 0.026, and a heterogeneity of 0.755. RGPs from phylogroup A are coloured in blue, from phylogroup B in yellow, and from phylogroup C in grey.

## Notes

https://gitlab.gwdg.de/botelho/pa_pangenome

### Competing Interest Statement

The authors have declared no competing interest.

### Summary of Updates

Revised text and figures, according to the reviewer's suggestions

