## Supplementary material for "Phylogroup-specific variation shapes the clustering of antimicrobial resistance genes and defence systems across regions of genome plasticity": Captions for supplementary tables

**Table S1**. General characteristics for the genomes discarded after quality control.

**Table S2**. General characteristics for the 2009 *P. aeruginosa* genomes used in this study.

**Table S3**. General characteristics for the 18 genomes that are part of the mPact strain panel described in this study.

**Table S4**. Total counts of gene families found across the pangenomes of different *P. aeruginosa* phylogroups.

**Table S5**. Comparison between the number of gene families for randomly subsampled 43 genomes from different phylogroups.

**Table S6**. Comparison between the number of gene families for randomly subsampled 100 genomes from phylogroups A and B.

**Table S7**. Presence/absence and classification of gene families across the different phylogroups.

**Table S8**. Functional annotation of the phylogroup-specific gene families.

**Table S9**. Virulence determinants found across the 2009 *P. aeruginosa* genomes from different phylogroups.

**Table S10**. CRISPR-Cas systems found across the 2009 *P. aeruginosa* genomes from different phylogroups.

**Table S11**. Antimicrobial resistance genes found across the 2009 *P. aeruginosa* genomes from different phylogroups.

**Table S12**. Defense systems found across RGPs and masked genomes (i.e., RGP-free genomes) from different phylogroups.

**Table S13**. ICEs and IMEs identified across the complete *P. aeruginosa* genomes from different phylogroups analysed in this study.

**Table S14**. General characteristics for the ICEs and IMEs identified in this study.

**Table S15**. General characteristics for the ICEs and IMEs network.
